# Unveiling potato cultivars with microbiome interactive traits for sustainable agricultural production

**DOI:** 10.1101/2024.08.21.609084

**Authors:** Tianci Zhao, Stefanie N. Vink, Xiu Jia, Alexander Erban, Stephanie Schaarschmidt, Joachim Kopka, Ellen Zuther, Krzysztof Treder, Dorota Michałowska, Rémy Guyoneaud, J. Theo M. Elzenga, Eléonore Attard, Joana Falcão Salles

## Abstract

Root traits significantly shape rhizosphere microbiomes, yet their interaction with microbes is often overlooked in plant breeding programs. Here, we propose that selecting modern cultivars based on microbiome interactive traits (MITs), such as root biomass, exudate patterns and the rhizosphere microbiome, can enhance agricultural sustainability by interacting effectively with soil microbiomes, which in turn, promotes plant growth and resistance to stress, thereby reducing reliance on synthetic crop protectants. Through a stepwise selection process (*in silico* and *in vitro*) that started with approximately 1000 potato genotypes, we chose 51 potato cultivars based on known phenotypical properties and distinct root exudate patterns. We conducted a greenhouse experiment to evaluate their capacity to interact with the soil microbiome and to assess their MITs. Our findings revealed that cultivars significantly influence plant growth, metabolite profiles, and rhizosphere fungal community composition. Moreover, we observed a positive correlation between microbial community diversity and root biomass. Additionally, leaf metabolites were correlated with rhizosphere bacterial composition, supporting the plant holobiont framework. Utilising z-scores, we aggregated all data related to plant growth, metabolomes, and microbiomes, creating a classification of 51 cultivars based on a gradient of MITs. By examining the distribution of low, medium, and high MITs, we identified a group of 11 potato cultivars suitable for further studies to assess their resilience and productivity under low-input production systems. This study provides an in-depth correlation between microbiome and several plant traits across 51 cultivars, offering tools to facilitate and expedite the incorporation of microbiome traits into breeding goals to support sustainable agriculture.

## 1 INTRODUCTION

Potato (*Solanum tuberosum* L.) cultivations rely heavily on conventional agricultural practices as a staple crop. Conventional management, which includes the widespread use of synthetic fertilisers and pesticides, is frequently used to enhance yields and protect crops from pests and diseases (Timsina, 2018). However, this dependency on synthetic compounds comes with substantial environmental issues. Overusing chemical inputs in conventional agriculture leads to water pollution, soil degradation, and soil microbial biomass and activity reduction, which are crucial in maintaining soil fertility and nutrient cycling (AL-Ani et al., 2019; Tripathi et al., 2020). Balancing high yields with care for our environment is a critical challenge in the breeding of crops.

In conventional breeding, plants are bred for traits such as high yield, disease resistance, tolerance to environmental stresses, and improved agronomic traits (Breseghello and Coelho, 2013). While pivotal in crop development, conventional breeding has inadvertently led to the dissociation between plant and soil microbiomes, negatively affecting beneficial plant-microbiome interactions (Spor et al., 2020). Comparisons between domesticated cultivars and their ancestral plants revealed that the rhizosphere microbiome of the latter exhibited higher complexity and connectivity (da Silva et al., 2023; Rossmann et al., 2020). However, relying solely on ancestral or wild plants may not effectively address the challenge of optimising plant-soil microbiome interactions in modern agricultural systems. Identifying specific genes associated with beneficial microbiomes in ancestral plants, with the aim of a subsequent transfer to modern crops (Clouse and Wagner, 2021), is complex due to environmental effects on gene expression (Raaijmakers and Kiers, 2022). Despite the reduced focus on plant-microbiome interactions during domestication (Wei and Jousset, 2017), multiple studies have demonstrated that inoculating modern crop cultivars with beneficial microbes can significantly enhance plant growth and stress resistance (Diagne et al., 2020; Fröhlich et al., 2012; Rodriguez et al., 2019; Tiwari et al., 2017). This suggests that modern cultivars retain the capacity for beneficial interactions with soil microbes, which can be leveraged or strengthened. Therefore, it is critical to identify gene markers associated with beneficial microbiomes in modern crops and apply this knowledge in real-world agricultural systems, particularly in the context of climate change.

The connection between root traits and plant productivity in crops has long been acknowledged, as roots are involved in resource acquisition, drought tolerance, soil exploration and other essential functions (Lynch, 1995; Ober et al., 2021). It is increasingly evident that our understanding of this connection should extend to how root traits shape the composition and functionality of the rhizosphere microbiome. While this aspect has received somewhat limited attention within plant breeding (Herms et al., 2022), many studies highlight the role of root traits in the rhizosphere microbiome. For instance, a study on maize revealed that wild maize had a significant impact on the structure of the microbial community, which was attributed to a high root-to-shoot biomass ratio (Szoboszlay et al., 2015). Plant root traits influence the soil microbial community by modulating the fungi-to-bacteria ratio, contributing to nutrient cycling (Wan et al., 2021). Moreover, a study involving beans (Pérez-Jaramillo et al., 2017) indicated that root architecture is linked to increased specific bacterial taxa, which could be connected to plant health. In addition to the root system, root exudates substantially impact the rhizosphere microbiome. As a survival strategy, plants secrete an abundance of compounds in their root exudates that can influence the diversity of the rhizosphere microbial community, promote the development of a more complex microbial network and enhance microbial carbon cycling, thereby facilitating the growth and survival of the host (Wang et al., 2022). A study on wheat has demonstrated that cultivars with a strong capacity for root exudates contribute to increased soil microbial diversity (Iannucci et al., 2021). Furthermore, the plant-associated microbial community is increasingly considered an extension of the plant phenotype (Bergelson et al., 2021; Whitham et al., 2003). Referred to as the plant’s “second genome”, plant-associated microbiome is strongly regulated by the plant’s genetic makeup (Turner et al., 2013). This suggests that a cultivar harbouring a more diverse community has a great potential to engage in beneficial interaction with microbes.

Here, we propose a method for identifying plant cultivars that foster beneficial interactions with the soil microbiome. We achieve this by assessing the microbiome interactive traits (MITs) of existing potato cultivars within the European potato germplasm bank. We hypothesise that root architecture and exudate patterns significantly influence the rhizosphere microbiome. By combining these traits, we propose a classification of cultivars with different MITs. Beginning with approximately 1000 potato genotypes, we employed a stepwise selection process, conducting in silico, *in vitro*, and greenhouse-based studies to examine the correlation between the plant traits of 51 potato cultivars and their rhizosphere microbial communities. We advocate for including MITs in breeding and the development of new crop cultivars. By prioritising these interactive traits, we can significantly enhance our understanding of plant-microbiome interactions while supporting microbiome-based innovation, which are essential for achieving sustainable agricultural practices.

## 2 MATERIALS AND METHODS

### 2.1 *In situ* analyses

In this project, we began our selection by screening late-season potato cultivars from a database of one thousand potato cultivars. Considering that disease-resistant plants are known to interact with beneficial microbes to bolster host immunity (Wille et al., 2019) and that these traits might be associated with their genetic background, we screened these cultivars from various resistance levels to numerous pathogens, especially those relating to potato-specific viruses (Table S1). The resistance scale was assessed using the method described by Michalak and Chrzanowska (2017). Detailed information on selected cultivars is available in The European Cultivated Potato Database (www.europotato.org). This selection resulted in a list of 148 to be used in the second selection round, which included a commercial cultivar, Desiree, as a reference.

### 2.2 *In vitro* experiment

For the second selection, the 148 cultivars including Desiree were evaluated according to the amount of dissolved organic carbon (DOC) in the root exudates of 6-week-old *in vitro* plants grown under sterile conditions (cf. section 2.1). Specifically, two-week-old potato plantlets grown from vegetative propagation in tissue culture at the Institute of Plant Breeding and Acclimation in Bonin (Bonin, Poland) were transplanted from agar tubes into 15 ml Eppendorf tubes, each containing 12 ml of sterile 0.5 x Hoagland solution. The agar was carefully removed from the roots of each plant, and to ensure aseptic conditions, sterile cotton wool was wrapped around the above-ground part of the plantlets, which were then gently placed into the 15 ml tubes. The roots were submerged in the Hoagland medium, while the above-ground part remained above it. The plantlets were allowed to acclimate to their new environment during a one-week growth period in a controlled climate chamber. The climate chamber maintained a temperature of 22 °C during the day and 18 °C at night, with a photoperiod of 16 hours of light and 8 hours of darkness. Upon completion of the initial week, the plantlets were transferred to new sterile 15 mL tubes, maintaining the 12 ml sterile 0.5 x Hoagland solution, as described in the previous step. To compensate for the liquid lost during the experiment, sterile water was added to maintain the volume at 12 ml. After the third week, the plants were harvested, and root and shoot length and dry weight were measured. Additionally, 3 ml aliquots of the growth medium were collected and frozen for DOC analysis. Moreover, 1 ml aliquots were freeze-dried for root exudate metabolite analysis (cf. section 2.4). The samples were immediately stored at -80°C until further analysis. The DOC content analysis was later processed in Helmholtz Zentrum München GmbH (Munich, Germany) (Data S1).

The data based on DOC content of the growth medium per root dry weight and shoot dry weight from the *in vitro* experiment revealed the variation among cultivars (Figure S1 and Table S2). We then selected a representative set of 50 cultivars plus the commercial cultivar, Desiree, representing the whole variation in DOC, for the third selection round in a greenhouse experiment.

### 2.3 Greenhouse experiment

The soil used in this experiment originated from a sugar beet field and was sieved with a 2 mm sieve. Before starting the experiment, six bulk soil samples were kept for physico-chemical analyses (Table S3) and DNA extraction. Two-week-old potato plantlets grown from vegetative propagation in tissue culture at the Institute of Plant Breeding and Acclimation in Bonin (Bonin, Poland) were transplanted from agar tubes into small pots containing soil (0.5 L) in the greenhouse. Before transplanting, the agar was carefully as previously described to ensure aseptic conditions. After two weeks of acclimatisation, the potato plants were transplanted into larger pots with the same soil (1.45 L). In the sixth week, the samples were taken as follows: Plant leaves, specifically the second or third leaf from the shoot top, were harvested for metabolomics analysis. The harvested leaves were snap-frozen in liquid nitrogen and then transferred to a freezer at -80 °C. The rhizosphere soil was obtained by gently brushing off adhering soil from the roots with a disposable toothbrush. Additionally, six bulk soil samples were collected from pots without plants for physicochemical analyses (Table S3) and DNA extraction. The soil samples for DNA analysis were frozen at -20 °C until DNA extraction the following day. Shoot height and root length were measured on the day of harvesting, and shoot and root dry weights were measured after drying at 60 °C for at least 48 hours.

### 2.4 Metabolite profiling of root exudates and leaf tissue

Metabolite fractions enriched for primary metabolites were profiled by gas chromatography and electron impact ionisation-time of flight mass spectrometry (GC/EI-TOF-MS) (Lisec et al., 2006; Erban et al., 2020). Root exudate samples, i.e. equal debris-free 1 mL volumes, of the *in vitro* experiment, cf. section 2.2, were freeze-dried directly without further sample preparation. A polar metabolite fraction was prepared from 50 mg snap-frozen leaf samples of the greenhouse experiment, cf. section 2.3. The frozen samples were extracted by a water-methanol-chloroform solvent mixture; a polar fraction was prepared from the extracts by water-induced liquid-liquid phase separation and dried in a speed-vacuum-concentrator, as described earlier (Erban et al., 2020). The dried fractions were subjected to methoxyamination and trimethylsilylation prior to GC/EI-TOF-MS analysis. ^13^C_6_ Sorbitol was added to all leaf samples before metabolite extraction (Erban et al., 2020). N-alkanes were added to each sample upon chemical derivatisation for subsequent retention index (RI) calibration (Erban et al., 2020). The GC/EI-TOF-MS chromatograms were obtained and baseline adjusted by ChromaTOF software (LECO Instrumente GmbH, Mönchengladbach, Germany) and background corrected using non-sample controls. Metabolite annotation was performed using TagFinder software (Luedemann et al., 2008), the NIST17 mass spectral database (U.S. Department of Commerce, Gaithersburg, USA), and the RI and mass spectral reference data of the Golm Metabolome Database, http://gmd.mpimp-golm.mpg.de/ (Hummel et al., 2010; Kopka et al., 2005). Compounds representing known contaminants and added internal standards were removed from further analysis. Metabolites absent from more than 75% of all analysed exudates or leaf tissue samples were excluded, resulting in 84 characterised leaf metabolites (Data S2) and 49 metabolites (Data S3) that were robustly present in root exudates.

Relative concentrations were normalised for further analyses to record the sample’s fresh weight and internal standard or exudate volume (Schaarschmidt et al., 2020). An ANOVA tool was used to perform a batch correction of the metabolite data sets for different measurement batches and the measurement sequence within batches (Lisec et al., 2011). All presented metabolite data are relative metabolite abundances. Three replicate samples per cultivar were analysed. The missing values were substituted by zero.

### 2.5 Soil sample sequencing and processing

DNA was extracted from 12 bulk soil samples and 153 rhizosphere samples with DNeasy PowerSoil Kit (Qiagen, Hilden, Germany) on 0.25 g of soil. DNA extraction followed the kit’s instructions except for the initial stage of bead beating, which was conducted with a FastPrep-24TM 5G Instrument at 6000 rpm/s for 40 s (MP Biomedicals, Santa Ana, USA). For sequencing of the bacterial community, the V4 region of the 16S rRNA gene was targeted using the primer sets 515F (5′-GTGCCAGCMGCCGCGGTAA-3′) and 806R (5′-GGACTACHVGGGTWTCTAAT-3′) (Caporaso et al., 2012, 2011). For the fungal community, the ITS2 region was sequenced with the primer sets 5.8SR (5’-TCGATGAAGAACGCAGCG-3’) and reverse primer ITS4 (5’-TCCTCCGCTTATTGATATGC-3’) (White et al., 1990). The Illumina MiSeq platform was used for paired-end sequencing (2 × 250bp for 16S, 2 × 300bp for ITS) at the PGTB (Genome Transcriptome Platform of Bordeaux, Cestas, France).

The 16S rRNA gene sequencing data were processed using a QIIME2 (version 2020.8) pipeline (Bolyen et al., 2019). Sequences were filtered, denoised, and dereplicated using the default setting of the Divisive Amplicon Denoising Algorithm (DADA2) plugin (Callahan et al., 2016). 16S rRNA taxonomic classification was performed using the q2-feature-classifier plugin (Bokulich et al., 2018) against the SILVA database (version 138) (Yilmaz et al., 2014). ITS sequencing data was processed using the PIPITS v. 2.4 pipeline (Gweon et al., 2015). In brief, the PEAR plugin was used to join read pairs (Zhang et al., 2014). The FASTX-Toolkit was utilised for quality filtering (Gordon and Hannon, 2010). The fungal-specific ITS2 region was extracted via ITSx (version 1.1b) (Bengtsson-Palme et al., 2013). The VSEARCH 2.13.3 plugin (Rognes et al., 2016) was used to dereplicate unique sequences, clustering to 97% sequence identity, and the UNITE Uchime reference dataset was used for chimera detection (Nilsson et al., 2015). Ultimately, the taxonomy was assigned with the RDP Classifier against the UNITE database (version 8.0) (Kõljalg et al., 2013).

### 2.6 Statistical Analysis

A one-way analysis of variance (ANOVA) was used to test variation in plant metabolite diversity, plant performance (shoot and root growth), and rhizosphere microbial alpha diversity (Data S2-S5). Post hoc comparisons were performed through Tukey’s honest significant differences or Duncan’s multiple range tests.

Analysis of soil microbial community structure (bacteria and fungi) was performed in R (version 4.2.0). Feature tables were rarefied at 14400 reads for bacterial 16S rRNA gene and 6361 for fungal ITS sequences, resulting in 8970 amplicon sequence variants (ASVs) for the bacterial community and 2006 operational taxonomic units (OTUs) for the fungal community after excluding non-microbial OTUs (Data S5). The rarefactions of feature tables were generated via ‘rarefy’ function in the package ‘vegan’ (Dixon, 2003). The same package was used to calculate the alpha diversity metrics for microbial communities and plant metabolites. Specifically, species richness indicates the number of unique species or metabolites observed, evenness describes how evenly the abundances of different species or metabolites are distributed, and the Shannon Diversity Index accounts for both the number of species/metabolites and their relative abundances (Jost, 2006; Wagner et al., 2018; Young and Schmidt, 2008).

The soil microbial community, root exudate and leaf metabolite compositions (Data S2, S3 and S5) were visualised through Principal Coordinate Analysis (PCoA) based on Bray-Curtis distance. This analysis used the ‘ape’ package in R (Paradis et al., 2004). PCoA scores per axis were averaged for each sample to illustrate the distribution of distinct cultivars in PCoA plots clearly. Permutational multivariate analysis of variance (PERMANOVA) was conducted using the ‘adonis’ function within the ‘vegan’ package to assess the impact of cultivars on soil microbial communities and metabolite profiles (Dixon, 2003).

Principal Component Analysis (PCA) based on the covariance matrix was performed with the R package ‘FactoMineR’ to reduce the dimensionality of the data and to visualise the distribution of plant cultivars (Lê et al., 2008). The variables included in the analysis were shoot and root length, shoot and root dry weight, metabolite diversity and composition of plant leaf tissue and root exudates (Data S4). Due to the significant variation in root exudate relative abundance observed in the *in vitro* experiment, median values were used for PCA plotting. Normalisation of plant data and metabolite richness was performed before analysis by calculating Z-scores. Subsequently, a PER-

MANOVA was conducted to assess the influence of variables on the distribution of 51 cultivars. The cultivars were classified into four functional groups based on their distribution across the first and second principal components (PC1 and PC2). The classification was done by dividing the PCA plot into four quadrants, each representing a distinct functional group of cultivars, reflecting differences in growth characteristics and metabolite profiles. The summarised group details can be found in Table S4.

To investigate the relationship between plant cultivars and rhizosphere microbiome composition, we categorised the 51 potato cultivars into four distinct functional groups. The category was determined by the distribution of cultivars and variables along the first two axes in the PCA plot.

The correlation between plant growth and related omics datasets (Data S4) was calculated using the Spearman correlation coefficient via the ‘corr.test()’ function from the ‘psych’ package in R (Revelle, 2024). The resulting p-values were adjusted using the False Discovery Rate (FDR) method to control for multiple comparisons and minimise false positives. We used the Mantel test with Spearman’s rank correlation to assess the correlation between microbial community compositions and metabolite profiles, utilising Bray-Curtis dissimilarity matrices. This analysis aimed to investigate the relationship between plant leaf metabolites and rhizosphere soil community.

To assess the overall MIT score of 51 potato cultivars, we calculated the average of the standardised scores (z scores) for various traits, including root length, root biomass, root-to-shoot biomass ratio, root exudate metabolites richness and Shannon diversity, as well as bacterial and fungal richness and Shannon diversity (Data S6)

Using the ‘ggsankey’ R package (Sjoberg, 2021), a Sankey plot was generated based on functional groups to summarise the selection of the 11 cultivars with the most potential for future research. The distribution of rhizosphere bacterial and fungal community compositions was illustrated for the 11 selected cultivars using Spearman correlation analysis. The first axis of the PCoA plot of bacterial and fungal communities served as the indicator of community composition (Data S4).

## 3 RESULTS

### 3.1 Plant growth

In the greenhouse experiment, plants from 51 potato cultivars were harvested from the soil in the sixth week, with plant growth varying among the cultivars. Analysis of variance (ANOVA) revealed that most of these cultivars’ growth patterns differed significantly according to their identity, with marked variation in both root biomass and root length (root biomass: *p* = 0.001, F = 2.26; root length: *p* = 0.003, F = 1.91; Figure 1). Specifically, the cultivars Inwestor, Salto and Szyper demonstrated higher root biomass than others. In addition, shoot length and shoot biomass varied significantly between cultivars (shoot length: *p* = 0.001, F = 6.80; shoot biomass: *p* = 0.009, F = 1.75; Figure S2). Here, the cultivars Astrid, Inwestor, and Tewadi cultivars had the highest shoot biomass. Regarding the root-to-shoot ratio based on length exhibited a significant response to distinct cultivars (*p* = 0.001; Figure S3) while the ratio of root-to-shoot dry weight did not show a substantial difference between the different cultivars (*p* = 0.36; Figure S3).

**Figure 1.**
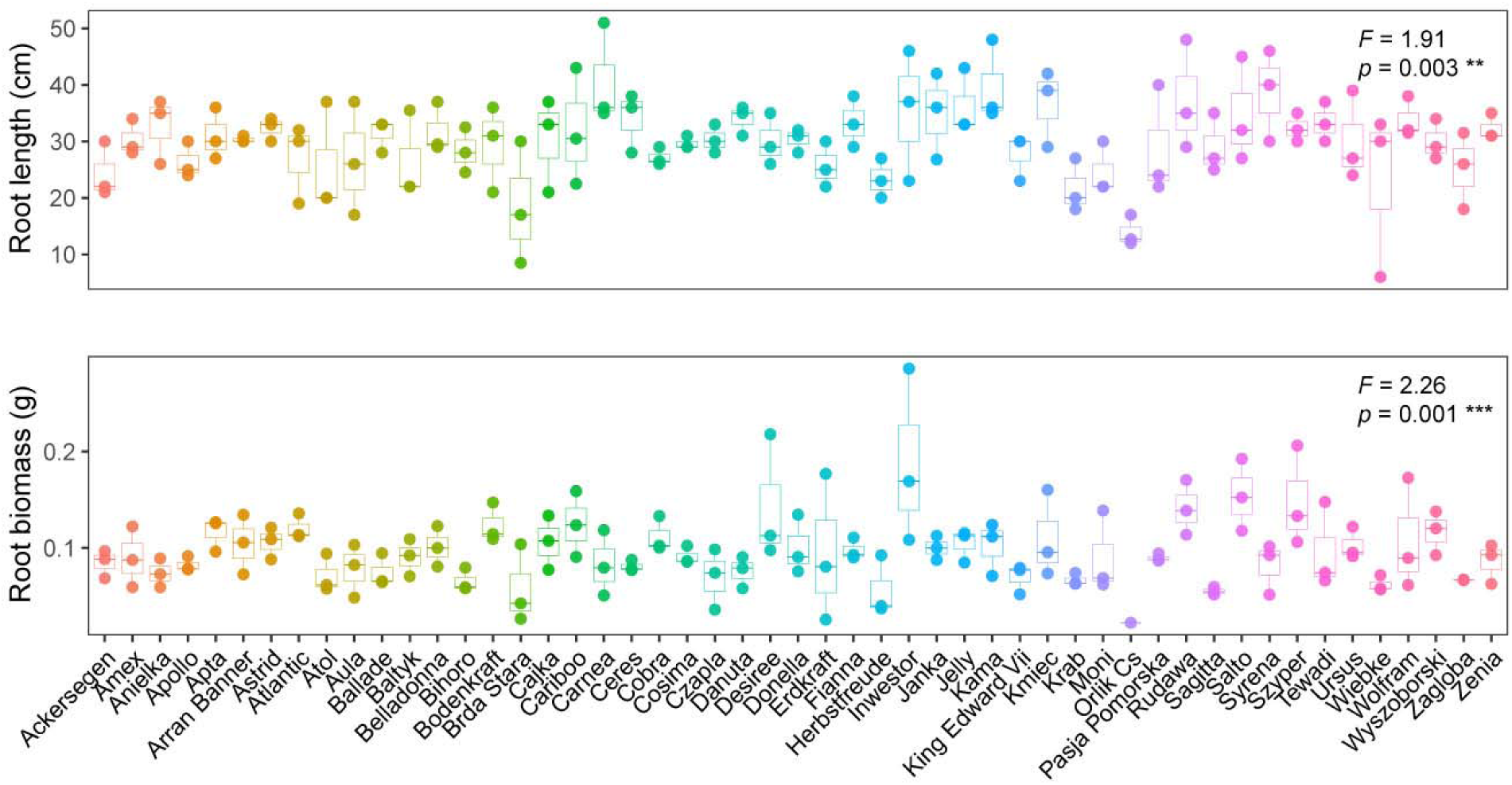
Root growth analysis of 51 potato cultivars. The upper panel displays root length, while the lower panel illustrates root dry weight. Each colour represents a distinct potato cultivar in alphabetical order. The upper right corner of each plot displays one-way ANOVA results, where the F-value explains the variation among different cultivars, and the *p*-value indicates the statistical relationship among cultivars. Significance levels are denoted as ** (*p* = 0.01) and *** (*p* = 0.001).

### 3.2 Plant leaf and root exudate metabolites

We further evaluated our selected 51 cultivars according to their root exudate metabolites (obtained under *in vitro* conditions) and leaf metabolites (obtained from the greenhouse experiment). To understand the variation in metabolites produced and released by the 51 potato cultivars, we conducted a Principal Coordinates Analysis (PCoA) and visualised the distribution of metabolites among different cultivars (Figure 2). The PERMANOVA results indicated significant differences in metabolite composition among cultivars for both root exudates and leaves (*p*_root exudate_ = 0.001 and *p*_leaf_ = 0.001; Figure 2). Overall, leaf metabolite composition was more strongly explained by plant cultivar identity than root exudate metabolites (*R*^2^_leaf metabolite_ = 0.68, and *R*^2^_root exudates_ = 0.58; Figure 2). The first two axes of the PCoA plot explained more than 34% of the variation for both metabolites.

**Figure 2.**
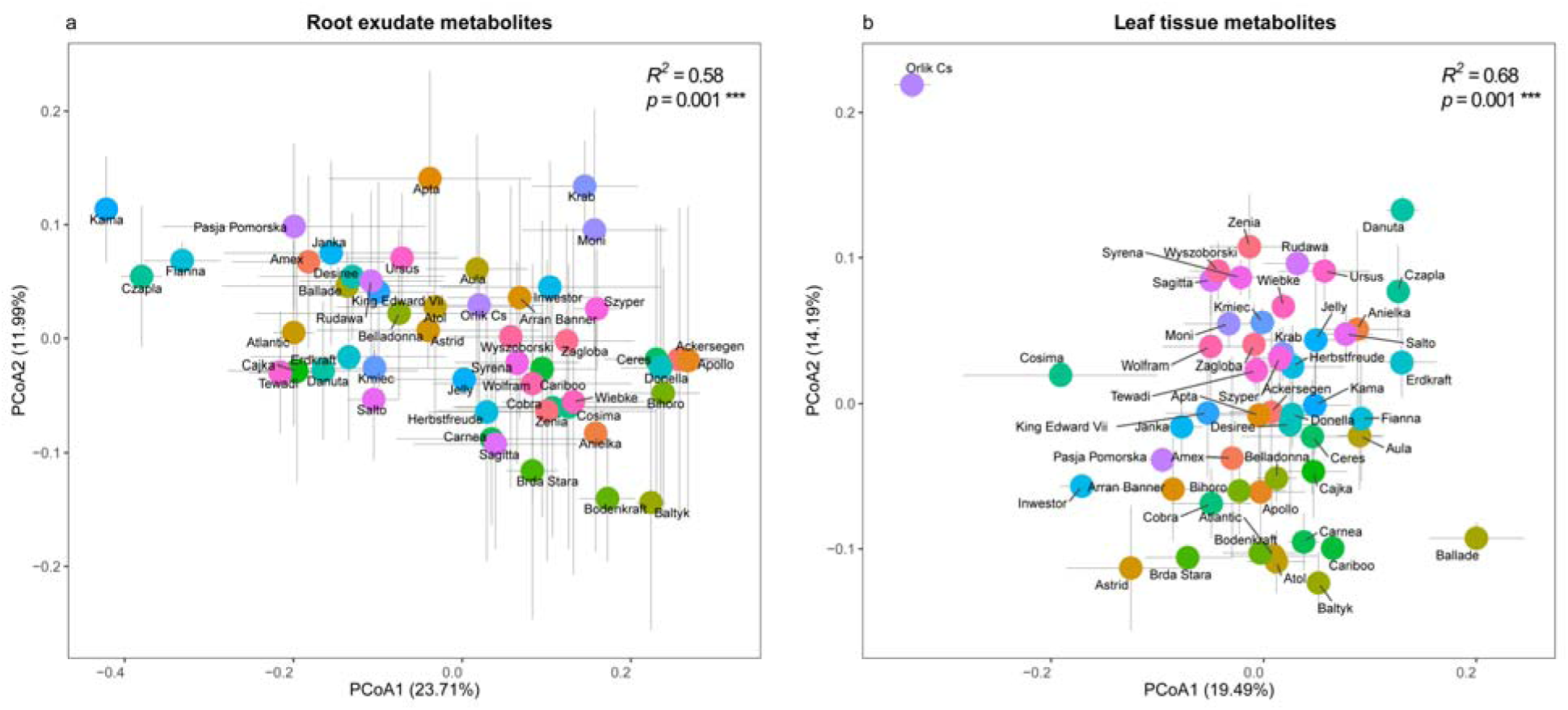
Distribution of metabolites in root exudates and leaves across 51 different cultivars. Principal Coordinates Analysis (PCoA) based on mean Bray-Curtis dissimilarity of metabolite composition of root exudates (a) and leaves (b). Distinct potato cultivars are represented by different colours, with the error bars for each cultivar displayed in grey. PERMANOVA results are shown in the upper right corner of each panel, indicating variations in metabolite composition among different cultivars. *R*² quantifies the explained variation, and *p*-values are derived from 9999 permutations; *** denotes statistically significant *p*-values (*p* = 0.001).

Although most cultivars tended to cluster together, several cultivars stood out, suggesting a different metabolite profiles. Specifically, Kama, Czapla and Fianna showed distinct root exudate metabolites compared to other cultivars, while Orlik Cs exhibited a unique leaf metabolite profile (Figure 2). To better illustrate the variance of remining cultivars, we remove these four cultivars with extreme metabolites values in Figure S4.

Furthermore, we found that the alpha diversity of plant metabolites was significantly affected by cultivars (Table S5). These results indicate that different potato cultivars exhibit distinct metabolites, contributing to variations in metabolite composition among cultivars.

### 3.3 Plant cultivar functional groups

To understand the distribution of plant growth and metabolite traits across our 51 selected cultivars, we used PCA to classify them into functional groups (Figure 3). These were based on data on plant cultivar growth characteristics, including shoot and root growth, leaf metabolites and root exudates. This approach allowed us to consider the overall metabolite composition, encompassing alpha and beta diversity, rather than focusing solely on metabolite richness and evenness. Our results revealed that the distribution of these traits differed significantly among different potato cultivars, as evidenced by the results of the PERMANOVA analysis (*p* = 0.001; Figure 3). Specifically, the first axis of the PCA biplot prominently reflected the impact of root exudate metabolite diversity and composition (Figure 3). In contrast, plant performance metrics and leaf metabolite composition mainly drove the second axis.

**Figure 3.**
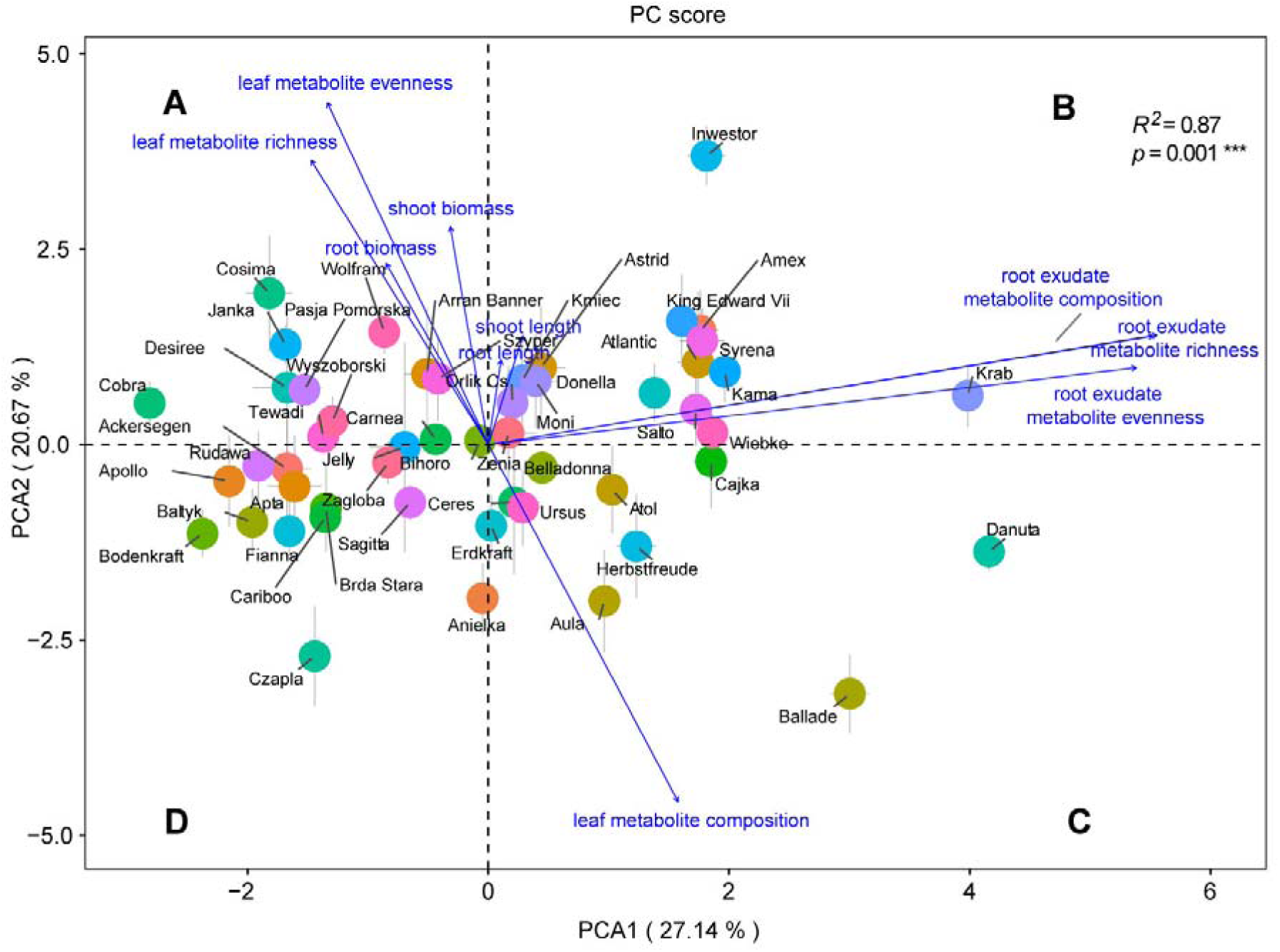
Distribution of 51 potato cultivars based on plant performance, root exudates and leaf metabolite profiles. Principal Component Analysis (PCA) based on the covariance matrix, highlighting variables impacting the distribution of potato cultivars. Distinct potato cultivars are represented by different colours, with the error bars for each cultivar displayed in grey. The result of PERMANOVA is shown in the upper right, elucidating the cultivar’s influence on the distribution. *R*² quantifies the explained variation, and p-values, derived from 9999 permutations, are denoted with *** for statistically significant results (*p* = 0.001). Letters A, B, C and D represent the four distinct functional groups.

Based on the distribution of cultivars and variables across the first two axes, the 51 potato cultivars were systematically categorised into four distinct functional groups according to their distribution in each of the four PCA quadrants (Figure 3). Group A featured cultivars exhibiting high leaf metabolite diversity and plant biomass, suggesting strong performance in leaf metabolic diversity and plant development. Group B included cultivars with high root exudate diversity and plant length, indicating a focus on below-ground metabolic processes and vertical growth of plants. Group C consisted of cultivars characterised by distinct leaf metabolite composition, as this variable primarily derived the cultivars distribution within this group. These cultivars may have a distinct composition of leaf metabolites compared to other groups, indicating specific leaf metabolic profiles that differentiate them from other cultivars. Group D comprised cultivars with low root exudate metabolite diversity and different metabolite compositions. The summarised group details and cultivars can be found in Table S4.

### 3.4 Soil microbial communities

The analysis of microbial communities in bulk and rhizosphere soil from the greenhouse experiment revealed a significant impact of plant presence on microbial communities. There was a significant difference in bacterial evenness between different soil compartments (*p* = 0.001; Figure S5b), whereas no significant effect on fungal richness and evenness (*p* > 0.05; Figure S5c,d) was observed. Bacterial and fungal community compositions responded to the presence of plants (Figure S6). Specifically, all bulk soil samples were found to cluster together in bacterial communities, separate from the rhizosphere samples (Figure S6a). Some potato cultivars’ rhizosphere fungal communities were grouped distinctly apart from the bulk soil (Figure S6b).

Comparison across the 51 potato cultivars revealed considerable variation and no significant effect of cultivar on bacterial and fungal alpha diversities, except for the cultivar effect on fungal evenness (*p* = 0.03; Figure S7). Plant cultivars showed no significant impact at the composition level but a trend towards influencing bacterial community distribution (*p* = 0.093 and *R*^2^ = 0.34; Figure 4a). However, cultivars exhibited significant variation in fungal community composition (*p* = 0.001 and *R*^2^ = 0.37; Figure 4b), indicating that cultivars can significantly influence the fungal community composition within their rhizosphere.

**Figure 4.**
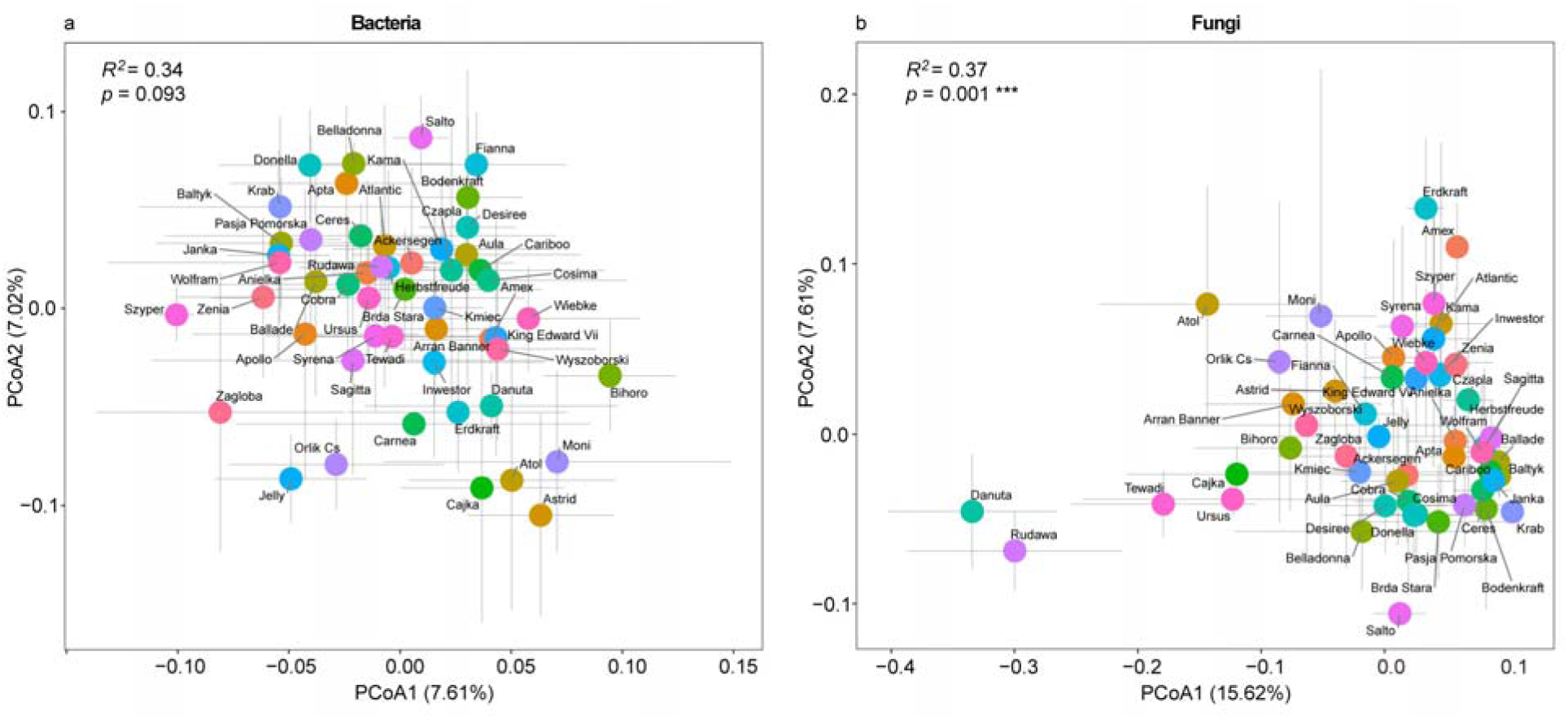
Rhizosphere microbial community composition. Principal Coordinates Analysis (PCoA) based on mean Bray-Curtis dissimilarity of community composition of bacteria (a) and fungi (b). Distinct potato cultivars are represented by different colours, with the error bars for each cultivar displayed in grey. PERMANOVA results in the upper left corner of each panel elucidate the influence of cultivars on community composition. *R*² quantifies the explained variation, and p-values are derived from 9999 permutations. The symbol *** denotes statistically significant p-values (*p* = 0.001).

Grouping the cultivars in the functional groups determined by MIT revealed no significant differences in the alpha diversity of bacterial and fungal communities between the different functional groups (*p* > 0.05; Figure S8). Similarly, the composition of rhizosphere microbial communities did not vary among the functional groups (*p*_bacteria_ = 0.49 and *p*_fungi_ = 0.14; Figure S9).

### 3.5 Correlation between plant traits, metabolites and rhizosphere microbiome

To evaluate how different plant traits relate to root-associated microbes and metabolite release, we correlated plant shoot biomass, root biomass and root exudate metabolites with leaf metabolites and rhizosphere microbial communities of the 51 potato cultivars. Positive correlations were observed between plant biomass and leaf metabolite richness. In contrast, negative correlations were identified between plant shoot growth and leaf metabolite composition (Figure 5). One possible explanation is that investment in growth represents a cost of metabolite synthesis but this needs to be proven by further experiments.

**Figure 5.**
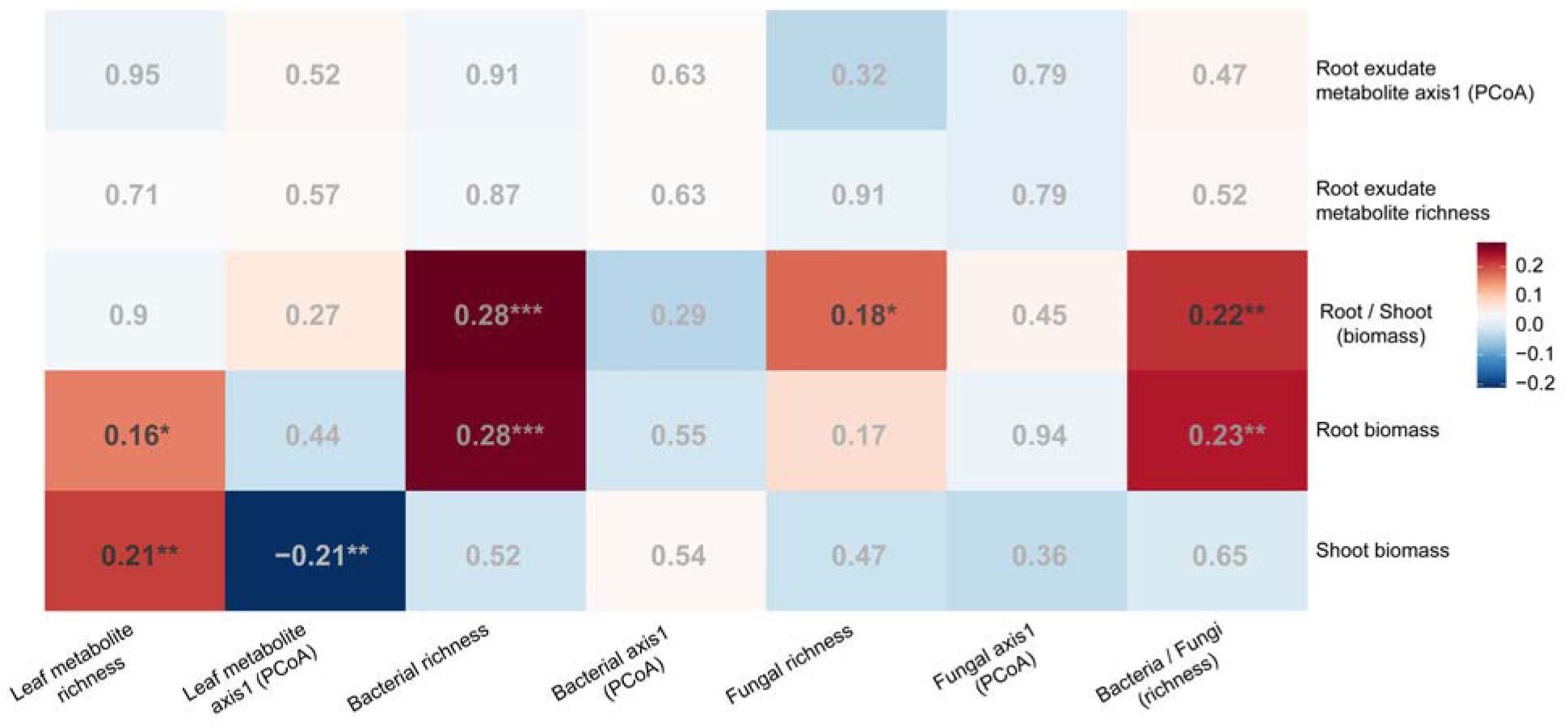
Correlation analysis between shoot biomass, MITs, leaf metabolite, and rhizosphere microbial community of the 51 potato cultivars. Correlations are based on the Spearman’s rank correlation coefficient. Dark-coloured boxes indicate a significant correlation (*p* < 0.05), with the colour intensity reflecting the strength of the correlation coefficient. Red represents a positive correlation, while blue represents a negative correlation. The numbers displayed within the boxes represent the correlation coefficient. Significance levels are denoted as follows: * (*p* < 0.05), ** (*p* < 0.01), and *** (*p* < 0.001).

Regarding the microbiome, we observed positive correlations between rhizosphere bacterial species richness, root biomass, and root-to-shoot biomass ratio (Figure 5). Similarly, rhizosphere fungal species richness positively correlated with the root-to-shoot biomass ratio. However, the composition of the rhizosphere microbial community showed no significant correlations with root growth. Notably, the bacteria-to-fungi species richness ratio (B/F) was positively associated with root and root-to-shoot biomass ratios. Additionally, we did not observe a pronounced correlation between root exudate metabolites and the rhizosphere microbial community (Figure 5).

We performed Mantel tests based on Bray-Curtis dissimilarity matrices to further investigate the relationship between plant leaf metabolites and soil microbial communities. Our analysis revealed a subtle yet statistically significant positive correlation between bacterial community compositions and leaf metabolite profiles (r = 0.10, *p* = 0.02). This suggests that plant cultivars with similar leaf metabolite profiles tend to harbour similar bacterial communities in the rhizosphere. In contrast, the Mantel test for fungi showed no significant correlation between fungal community compositions and leaf metabolite profiles (r = 0.04, *p* = 0.22).

When examining the correlations between plant root biomass and rhizosphere bacterial community diversity across different functional groups, functional groups A and D showed significant positive correlations (*p* < 0.05; Figure S10a,b). Functional groups B and C show no significant correlation between these variables. There were no significant correlations between root biomass and fungal community diversity (*p* > 0.05; Figure S10c, d) across functional groups.

### 3.6 Classification of cultivars based on MITs

To evaluate the overall MIT scores of different cultivars, we calculated the average of the standardised scores (z scores) for various traits, including root length, root biomass, root-to-shoot biomass ratio, root exudate metabolites richness and Shannon diversity, as well as bacterial and fungal richness and Shannon diversity. Based on these MIT z-score values, the 51 cultivars were categorised into either high, middle, or low MIT levels (Figure 6a; Table S6). Furthermore, we illustrated the distribution of the rhizosphere community composition of the 51 cultivars in Figure 6b, which shows a substantial separation in bacterial and fungal compositions among the MIT-selected cultivars.

**Figure 6.**
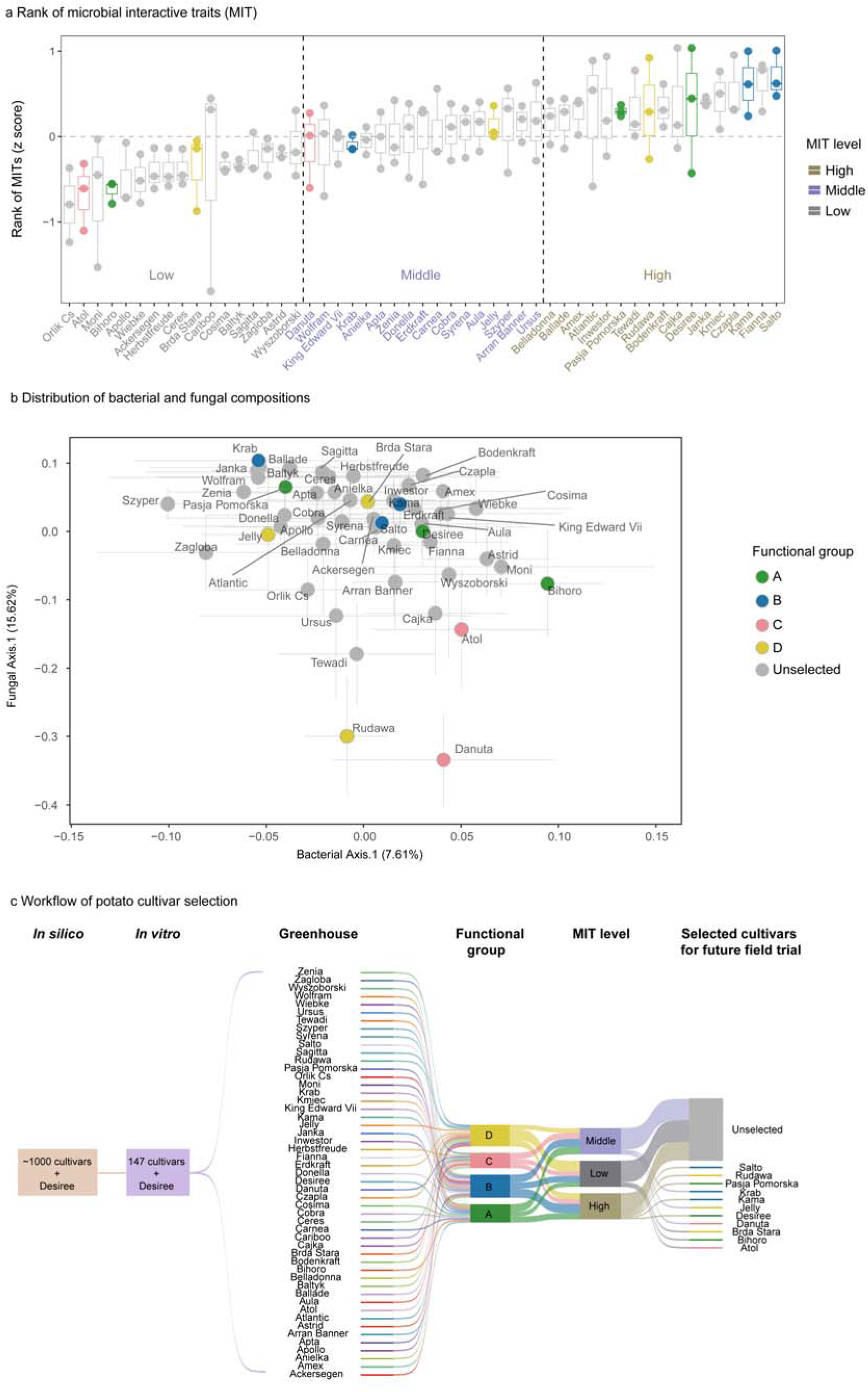
Rank of 51 potato cultivars based on microbial interactive trait (MIT) z-scores (a). Cultivars were ranked according to the standardised (z-score) values of multiple MIT-related parameters, including root length, root biomass, root-to-shoot biomass ratio, richness and Shannon diversity of root exudate metabolites, as well as bacterial and fungal richness and Shannon diversity. Based on their overall z-scores, cultivars were grouped into high, middle, and low MIT levels. Distribution of rhizosphere bacteria and fungi community composition through Spearman correlation analysis (b). The first axis of principal coordinate analysis (PCoA1) of microbial community beta-diversity (bacteria and fungi) serves as the indicator of community composition. The error bars for each cultivar are displayed in grey. Workflow of potato cultivar selection (c). This Sankey plot outlines the selection process of potato cultivars based on functional group classification and MIT levels. The colour of selected cultivars corresponds to functional groups. The cultivars that were not selected are depicted in grey.

### 3.7 Summary of the selection procedure

From the different functional groups (Figure 3) and MIT levels, we suggested 10 cultivars plus Desiree for future studies (Figure 6c, Table S6). The stepwise selection workflow and the selected cultivars are visualised in Figure 6c. We initially selected 148 potato cultivars, including the commercial cultivar Desiree as a reference, from a pool of a thousand based on their resistance to pathogens *in silico*. These 148 cultivars were then further grown *in vitro*, from which 51 were selected based on DOC content in root exudates for a subsequent greenhouse experiment. For these 51 cultivars, plant growth, leaf metabolites and root exudate metabolites were used to classify them into four functional groups with distinct growth traits. MIT scores allowed us to identify ten representative cultivars with diverse MIT levels across different functional groups, which should undergo further exploration in real-world conditions to evaluate their potential for interacting with beneficial soil microbiomes.

We conducted a comprehensive characterisation of the MIT-selected cultivars (Figure S11). The one-way ANOVA results revealed variations among the selected 11 cultivars (10 + Desiree) in terms of plant metabolites, root development, and rhizosphere microbial community richness. Specifically, Desiree and Pasja Pomorska from group A demonstrated higher leaf metabolite richness and microbial diversity (Figure S11a, c, d, g, h). In contrast, in group B, Krab and Salto showed superior microbial alpha diversity performance (Figure S11c, d, g, h). In comparison, Danuta from group D exhibited lower plant metabolite richness, root growth, and bacterial diversity (Figure S11a-c, e-g). Regarding beta diversity, there were no significant cultivar effects on metabolite or microbial profiles, except for root exudate metabolite composition (Figure S11i-l).

## 4 DISCUSSION

Conventional breeding primarily harnesses genetic variation to achieve the desired traits of plants. However, the impact of a cultivar’s genetic information on associated microbial organisms has yet to receive much consideration, particularly those associated with the rhizosphere. Here, we emphasise that modern breeding should consider the soil microbiome as a strategy to reduce its environmental footprint. Understanding the importance of microbiome interactive traits (MITs), such as root traits and exudates, is vital to comprehending how plants and microbiomes interact. Below, we discuss our findings in the context of the correlation between the selected MITs, their potential contribution to sustainable agriculture and additional traits that could be considered in future studies. The integration of this knowledge will contribute to informing the breeding process, providing valuable insights for developing new microbiome-based cultivars.

### 4.1 Correlation between plant traits, metabolites and rhizosphere microbiome

Our findings underscore the significant role of plant genetics in shaping both root exudate and leaf metabolite profiles. We observed substantial cultivar-dependent variation in root exudate metabolite composition consistent with previous studies. This genetic influence also extends to leaf metabolites, aligning with earlier research on rice (Schaarschmidt et al., 2020). These results collectively demonstrate that the genetic makeup of different plant cultivars significantly influences the metabolite composition of different plant tissues.

We observed a positive correlation between leaf metabolite richness and plant biomass, a relationship that can be explained through considerations of resource availability and plant growth strategies. Environmental conditions and functional traits influence biomass allocation among plant organs, such as leaves and roots, influencing many growth processes (Mensah et al., 2016; Poorter et al., 2012). In optimal environmental conditions, biomass allocation (primarily above-ground to compete for light) can enhance the diversity of leaf metabolites instead of root metabolites (Chapin et al., 2005). The crucial role of plant secondary metabolites in host defence may also explain this correlation. These compounds protect hosts against herbivores, pathogens, and (biotic) stresses (Anjali et al., 2023; Divekar et al., 2022; Yadav et al., 2021; Zaynab et al., 2018). Plants with higher biomass may invest more in defence mechanisms, producing a wider range of leaf metabolites for protection. This establishes positive feedback loops, wherein increased biomass boosts photosynthetic activity and energy production, consequently supporting the synthesis of more diverse metabolites.

Previous studies have demonstrated that plant root traits are crucial in regulating rhizosphere microbial communities (Szoboszlay et al., 2015; Wan et al., 2021). This relationship is further supported by Eisenhauer et al. (2017), who showed that microbial diversity increases with increasing root biomass and exudate amount. Our results align with these findings, revealing positive correlations between plant root biomass and root-to-shoot biomass ratio, rhizosphere bacterial diversity, and bacteria-to-fungi species richness ratio (B/F). A more diverse microbial community, in turn, can enhance soil nutrient cycling and availability (Jiao et al., 2021), potentially promoting plant growth.

Although root exudates significantly influence bacterial and fungal communities in the rhizosphere (Hartmann et al., 2009), our study did not find a significant correlation between root exudate metabolites and the rhizosphere microbial community. This unexpected result may be because we collected root exudate metabolites from 51 potato cultivars under *in vitro* conditions, while the microbial community data were obtained from a greenhouse experiment. The differences in substrates between these environments may have influenced the root exudate profiles, leading to variances that could explain the lack of correlation.

Given the complexity of collecting root exudates from soil plants, we collected leaf metabolites in our current experiment to explore the relationship between plant metabolites and the soil microbial community. The marginal but positive correlation observed between leaf metabolites and the rhizosphere bacterial community suggests a potential influence of leaf metabolites on the composition of rhizosphere bacteria. This finding indicates a possible link between above-ground plant tissues and below-ground microbial communities, aligning with the holobiont concept, in which plants and their associated microorganisms are viewed as a holistic ecological unit (Vandenkoornhuyse et al., 2015).

Recent studies further support this interconnection. Korenblum et al. (2020) demonstrated that the composition of the rhizosphere microbial community affects the metabolomes and transcriptomes of tomatoes’ leaves and roots. Earlier research showed that *Bacillus* can influence photosynthesis, leaf growth, and overall plant phenotypes by producing phytohormones or volatile organic compounds (Pang et al., 2021), potentially impacting leaf metabolomes. These findings collectively emphasise the concept of metabolites as primary mediators regulating plant-microbiome interactions within the holobiont framework (Carper et al., 2022). As described, the transport of leaf-produced metabolites to the roots via the phloem (Broussard et al., 2023) suggests a potential mechanism for how above-ground metabolites might influence root exudate patterns and, consequently, the rhizosphere microbiome. However, it’s important to note that our study only demonstrates correlation, not causation. The complex interactions between leaf metabolites, root exudates, and microbial communities require further investigation to elucidate the holobiont concept fully.

### 4.2 Enhancing plant-microbiome interactions for sustainable agriculture

The soil microbiome promotes plant growth by promoting carbon, nutrient, and phosphorus cycling (Hartmann and Six, 2022). It also contributes to plant resistance by producing hormones that protect against abiotic and biotic stress (Eichmann et al., 2021). Despite its critical functions, the soil microbiome has received limited attention in conventional breeding (Mitter et al., 2019; Wei and Jousset, 2017). Compounding this issue is the fact that conventional agricultural management not only negatively affects the environment but also substantially impacts the soil microbiome (Longepierre et al., 2021). This dual impact increases the decoupling between plants and soil microbiomes (Huang et al., 2019; Spor et al., 2020).

Studies addressing the role of plants in regulating their associated microbiome remain relatively limited (Wei and Jousset, 2017). More precisely, plant roots, serving as the primary interface for interaction with soil microbes, are underexplored in plant breeding (Herms et al., 2022; Reinhold-Hurek et al., 2015). Here, we consider morphological root characteristics and root exudate metabolites as MITs to explore their interaction with the rhizosphere microbiome. We aim to supply a strategy for breeding that considers plant-associated microbiota.

Indeed, the positive correlation between root growth and rhizosphere microbial diversity indicates that MITs can aid in identifying plant cultivars with the potential to interact effectively with root-associated microbiomes. Cultivars exhibiting high MITs are likely to harbour a more diverse rhizosphere microbiome, which in turn can lead to enhanced plant growth through the support of beneficial microbial interactions. Identifying the genes associated with beneficial microbiomes in modern cultivars and using them in selective breeding efforts to achieve microbial-assisted cultivars can serve as a new plant breeding strategy. This approach represents a promising avenue for sustainable agriculture, as it harnesses the power of beneficial microorganisms to improve crop performance while reducing the need for chemical inputs.

### 4.3 Integrating additional root traits and phyllosphere microbiome in future studies

We suggest expanding future research beyond the MITs examined in this study to include a broader range of root phenotypic traits. While the current study focuses on root biomass and length, future investigations should include root diameter, surface area, and root type. Although less studied, evidence suggests that fine roots with smaller diameters have larger surface areas, potentially recruiting a greater diversity and abundance of microbes through enhanced nutrient and metabolite exchange (Saleem et al., 2018; Wan et al., 2021). Pérez-Jaramillo et al. (2017) linked root types (thin or thick) to specific bacterial phyla, highlighting the importance of root morphology in shaping microbial communities. Additionally, the spatial distribution of microbial communities along the root should be considered. Kawasaki et al. (2016) observed that the functional genes detected in microorganisms near the root tip were distinct from those isolated near the root base. Collectively, these various root phenotypic traits should be considered to improve plant-microbiome interactions.

In addition to expanding our focus on root phenotypic traits, we propose incorporating the phyllosphere microbiome into future studies. The phyllosphere microbiome, which includes microorganisms inhabiting the above-ground parts of plants, plays a crucial role in plant health and function (Thapa and Prasanna, 2018; Vorholt, 2012). Previous studies have demonstrated that host genotypes significantly influence the composition of phyllosphere microbial communities (Bodenhausen et al., 2014; Thapa et al., 2017). The phyllosphere microbiome is involved in nitrogen fixation (Abadi et al., 2021), enhancing stress tolerance (Etemadi et al., 2018; Stone et al., 2018), and suppressing plant diseases (Fan et al., 2019; Das et al., 2023). Additionally, they can regulate plant growth through the production of plant hormones (Stone et al., 2018). These diverse functions highlight the importance of the phyllosphere microbiome in plant health and productivity.

By considering the phyllosphere microbiome together with the rhizosphere microbiome and plant metabolites, we can establish a more comprehensive understanding of the plant holobiont. This approach will allow us to bridge the gap between the plant’s above- and below-ground components. By harnessing the functions of phyllosphere and rhizosphere microbiomes, we may enhance crop yields, improve plant resilience, and reduce reliance on chemical inputs, ultimately contributing to more sustainable agricultural systems.

## 5 CONCLUSION

This study underscores the significant impact of plant cultivars on leaf metabolites and root exudate metabolites. We also observe a positive correlation between leaf metabolites and rhizosphere bacterial community; further studies are needed to verify the causation and to involve root exudates to expand our knowledge of the holobiont framework. We systematically selected potato cultivars to identify those with diverse microbiome interactive traits (MITs). We lay the foundation for further studies to evaluate the performance of MIT-selected cultivars in the real world. This is needed to provide a promising strategy for future breeding programs, including identifying gene markers associated with a beneficial microbiome and utilising these genes to increase plant-microbiome interactions. This breeding strategy could promote host growth while reducing the reliance on synthetic chemicals in conventional agriculture. Finally, we suggest integrating additional root phenotypic traits and the phyllosphere microbiome in future studies to establish a more comprehensive understanding of the plant holobiont, which can benefit plant-microbiome interactions.

## Data Availability Statement

The raw sequencing data are available in the National Center for Biotechnology Information (NCBI) Sequence Read Archive (SRA) under the accession number PRJNA1211026. The metadata and datasets used for the bioinformatic analyses are available at the following link: https://github.com/tianci-zhao/potatoMETAbiome-Greenhouse-Experiment.

## ACKNOWLEDGEMENTS

This study was funded by ERA-NET Cofund SusCrop project potatoMETAbiome, which is supported by the European Union’s Horizon 2020 research and innovation program (grant agreement No 771134; French National Agency, grant number-ANR-18-SUSC-0001) and part of the Joint Programming Initiative on Agriculture, Food Security and Climate Change (FACCE-JPI). We thank Jolanda K. Brons, Jan Veldsink, Xipeng Liu, Pina Brinker, Panji Cahya Mawarda, Edisa Garcia Hernandez, Georgia Voulgari, and Florian Fischer for their help in the lab. We thank Michael Schloter, Benoit Renaud Martins, Viviane Radl, and Gudrun Hufnagel from Helmholtz Zentrum München GmbH for their contributions to DOC samples processing. We thank Ines Fehrle and Jessica Alpers for their excellent assistance with the GC-MS measurements. We want to thank Marie Simonin and Ulisses Nunes da Rocha for their insightful discussion on data analysis. T.Z. was supported by a scholarship from the China Scholarship Council (CSC) and the University of Groningen scholarship program. We thank PGTB for sequencing (Genome Transcriptome Platform of Bordeaux). We thank the Center for Information Technology at the University of Groningen for providing access to the Hábrók high-performance computing cluster.

## Author contributions

J.F.S., S.N.V., J.T.M.E., A.E., and E.A. designed the experiment; T.Z. and S.N.V. performed the experiment, laboratory work and data analysis; X.J. participated in data analysis; E.A. and R.G. contributed to plant sampling; D.M. and K.T. provided the potato materials; A.E., S.S., J.K., and E.Z. measured and analysed plant metabolites. T.Z. drafted the manuscript, with all authors contributing to its modification and approval of the final version.

## Supporting information

**Figure S1.** Root dry weight and root-to-shoot ratio of 51 potato cultivars *in vitro* experiment. Each colour represents a distinct potato cultivar. The upper left corner of each plot displays one-way ANOVA results, where the F-value explains the variation among different cultivars, and the *p*-value indicates the statistical relationship among cultivars.

**Figure S2.** Shoot growth analysis of 51 potato cultivars. The upper panel displays the shoot length, while the lower panel illustrates the shoot dry weight. Each colour represents a distinct potato cultivar. The upper right corner of each plot displays one-way ANOVA results, where the F-value explains the variation among different cultivars, and the *p*-value indicates the statistical relationship among cultivars.

**Figure S3.** Root-to-shoot ratio analysis of 51 potato cultivars. The upper panel displays the root-to-shoot length ratio, while the lower panel illustrates the root-to-shoot dry weight ratio. Each colour represents a distinct potato cultivar. The upper right corner of each plot displays one-way ANOVA results, where the F-value explains the variation among different cultivars, and the *p*-value indicates the statistical relationship among cultivars.

**Figure S4.** Distribution of metabolites in root exudates and leaves across 47 different cultivars. Four outlier cultivars (Kama, Czapla, Fianna, Orlik Cs) were excluded to better illustrate the distribution of distinct cultivars. A Principal Coordinates Analysis (PCoA) based on Bray-Curtis dissimilarity was performed to visualise the composition. The metabolite dissimilarities of root exudates and leaf are depicted separately on the left and right. Distinct potato cultivars are represented by different colours, with the error bars for each cultivar displayed in grey. PERMANOVA (Adonis) results in the upper right corner of each panel elucidate the influence of cultivars on metabolite composition. *R*² quantifies the explained variation, and *p*-values are derived from 9999 permutations. The symbol *** denotes statistically significant *p*-values (*p* = 0.001).

**Figure S5.** The alpha-diversity of the bacterial (a,b) and fungal (c,d) communities in bulk and rhizosphere soil, displayed by species richness and evenness. Different colours represent bulk samples from the beginning of the experiment (Bulk_D0), at harvest (Bulk_D36), and rhizosphere samples of 51 cultivars. The lower right corner of each plot displays one-way ANOVA results, where the F-value explains the variation among different soil compartments (bulk and rhizosphere), and the *p*-value indicates the statistical relationship.

**Figure S6.** The microbial community composition in bulk and rhizosphere soil. Principal Coordinates Analysis (PCoA) based on Bray-Curtis dissimilarity was performed to visualise the community composition of bacteria (a) and fungi (b). Different shapes represent different soil compartments (bulk and rhizosphere). Different colours represent bulk samples from the beginning of the experiment (Bulk_D0), at harvest (Bulk_D36), and different potato cultivars, with the error bars for each cultivar displayed in grey. PERMANOVA results in the lower left corner of each panel elucidate the influence of soil compartments on community composition. *R*² quantifies the explained variation, and *p*-values are derived from 9999 permutations.

**Figure S7.** Rhizosphere microbial community evenness of 51 potato cultivars. The upper panel is bacterial evenness, and the lower panel is fungal evenness. Different colours indicate different cultivars. The lower right corner of each plot displays one-way ANOVA results, where the F-value explains the variation among different cultivars, and the *p*-value indicates the statistical relationship among cultivars.

**Figure S8.** Rhizosphere microbial community species richness. The upper panel is bacterial richness, and the lower panel is fungal richness. Different colours indicate different cultivar functional groups. The “ns” indicates no significant influence of groups on microbial alpha diversity (one-way ANOVA).

**Figure S9.** Rhizosphere microbial community composition. Principal Coordinates Analysis (PCoA) based on Bray-Curtis dissimilarity was performed to visualise the community dissimilarities of bacteria (a) and fungi (b). Distinct potato cultivar functional groups are represented by different colours, with the error bars for each cultivar displayed in grey. PERMANOVA results in the upper right corner of each panel elucidate the influence of groups on community composition. *R*² quantifies the explained variation, and *p*-values are derived from 9999 permutations.

**Figure S10.** Correlation between plant root biomass and rhizosphere bacterial (a,b) and fungal (c,d)alpha diversities (species richness and Shannon diversity) across different functional groups. Displayed by functional groups A, B, C and D. The Spearman correlation assessed the relationship, with *R*^2^ indicating the strength of the correlation. Y is the regression equation, and a *p*-value < 0.05 represents a significant correlation between the variables.

**Figure S11.** Characterisation of selected cultivars. Selected potato cultivars from different functional groups are represented by different colours, with the error bars for each cultivar displayed in grey. Letters in the upper two panels indicate significant differences across cultivars (Duncan post hoc test). In the lower right corner of the last panel plots, PERMANOVA results elucidate the influence of cultivars on community composition. *R*² quantifies the explained variation, and p-values are derived from 9999 permutations. A significance level is denoted as *** (*p* < 0.001).

**Table S1.** Background information on 148 selected cultivars for the *in vitro* experiment.

**Table S2.** Plant growth data of 148 potato cultivars from *in vitro* experiment.

**Table S3.** Soil physicochemical characteristics at the beginning and end of greenhouse experiment.

**Table S4.** Functional groups of potato cultivars categorised based on plant growth and metabolite pro-files.

**Table S5.** The one-way analysis of variance (ANOVA) shows the influence of cultivar on the plant metabolite alpha diversity.

**Data S1.** Dissolved organic carbon content of root exudates from *in vitro* experiment

**Data S2.** Leaf tissue metabolites data in greenhouse experiment

**Data S3.** Root exudate metabolites data from *in vitro* experiment

**Data S4.** Plant performance in greenhouse experiment

**Data S5.** Rhizosphere microbial feature tables

**Data S6.** MIT z scores data

