## Supplement for "Unveiling potato cultivars with microbiome interactive traits for sustainable agricultural production"

**SUPPORTING INFORMATION**

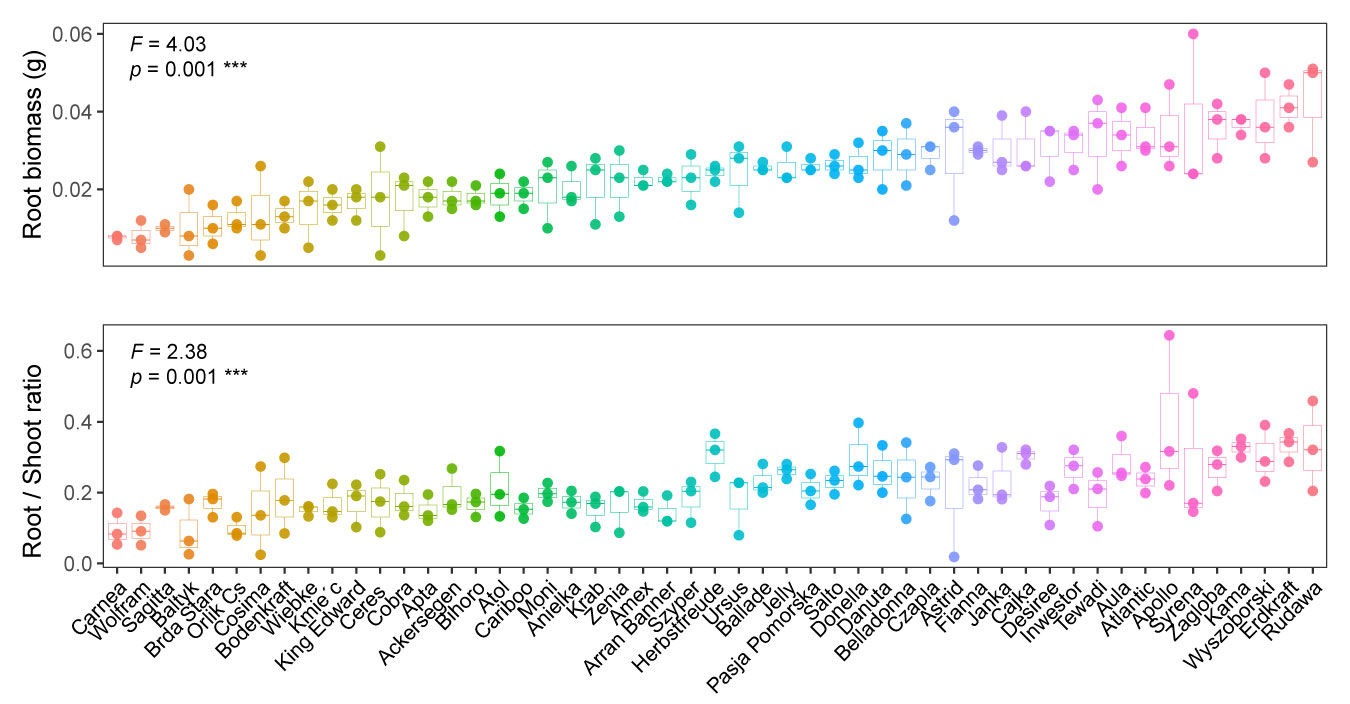
**Figure S1.** Root dry weight and root-to-shoot ratio of 51 potato cultivars *in vitro* experiment. Each colour represents a distinct potato cultivar. The upper left corner of each plot displays one-way ANOVA results, where the F-value explains the variation among different cultivars, and the *p*-value indicates the statistical relationship among cultivars. Significance levels are denoted as *** (*p* = 0.001).

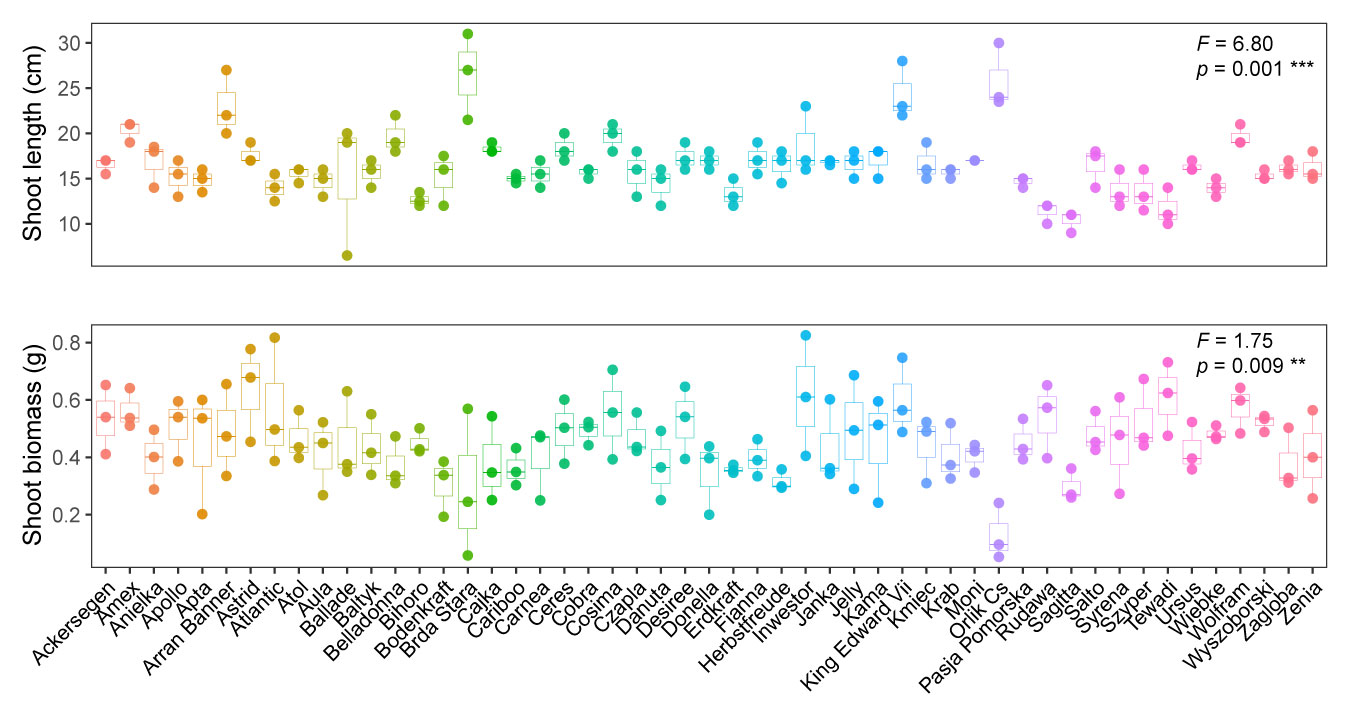
**Figure S2.** Shoot growth analysis of 51 potato cultivars in greenhouse condition. The upper panel displays the shoot length, while the lower panel illustrates the shoot dry weight. Each colour represents a distinct potato cultivar. The upper right corner of each plot displays one-way ANOVA results, where the F-value explains the variation among different cultivars, and the *p*-value indicates the statistical relationship among cultivars. Significance levels are denoted as ** (*p* = 0.01) and *** (*p* = 0.001).

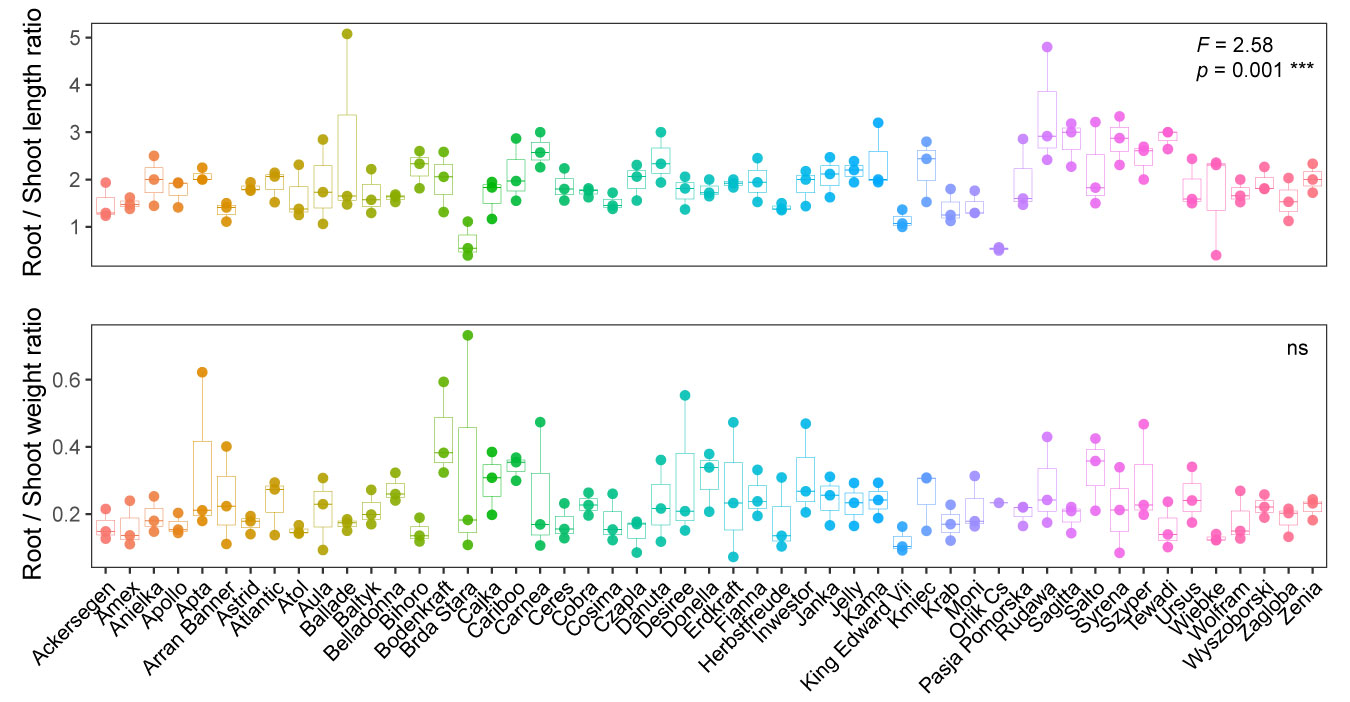
**Figure S3.** Root-to-shoot ratio analysis of 51 potato cultivars growth in greenhouse condition. The upper panel displays the root-to-shoot length ratio, while the lower panel illustrates the root-to-shoot dry weight ratio. Each colour represents a distinct potato cultivar. The upper right corner of each plot displays one-way ANOVA results, where the F-value explains the variation among different cultivars, and the *p*-value indicates the statistical relationship among cultivars. Significance levels are denoted as *** (*p* = 0.001). The “ns” indicates no significant difference.

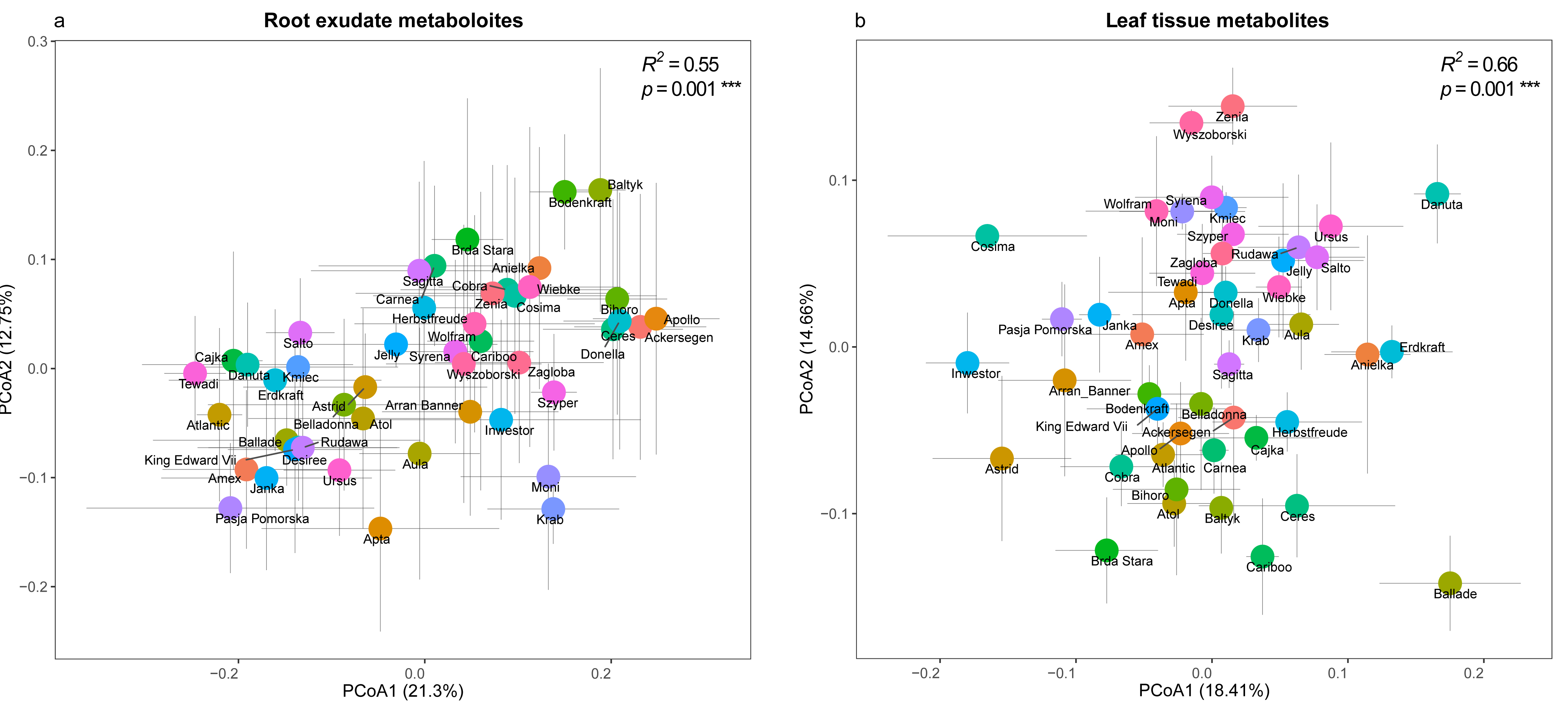
**Figure S4.** Distribution of metabolites in root exudates and leaves across 47 different cultivars. Four outlier cultivars (Kama, Czapla, Fianna, Orlik Cs) were excluded to better illustrate the distribution of distinct cultivars. A Principal Coordinates Analysis (PCoA) based on Bray-Curtis dissimilarity was performed to visualise the composition. The metabolite dissimilarities of root exudates and leaf are depicted separately on the left and right. Distinct potato cultivars are represented by different colours, with the error bars for each cultivar displayed in grey. PERMANOVA (Adonis) results in the upper right corner of each panel elucidate the influence of cultivars on metabolite composition. *R*² quantifies the explained variation, and *p*-values are derived from 9999 permutations. The symbol *** denotes statistically significant *p*-values (*p* = 0.001).

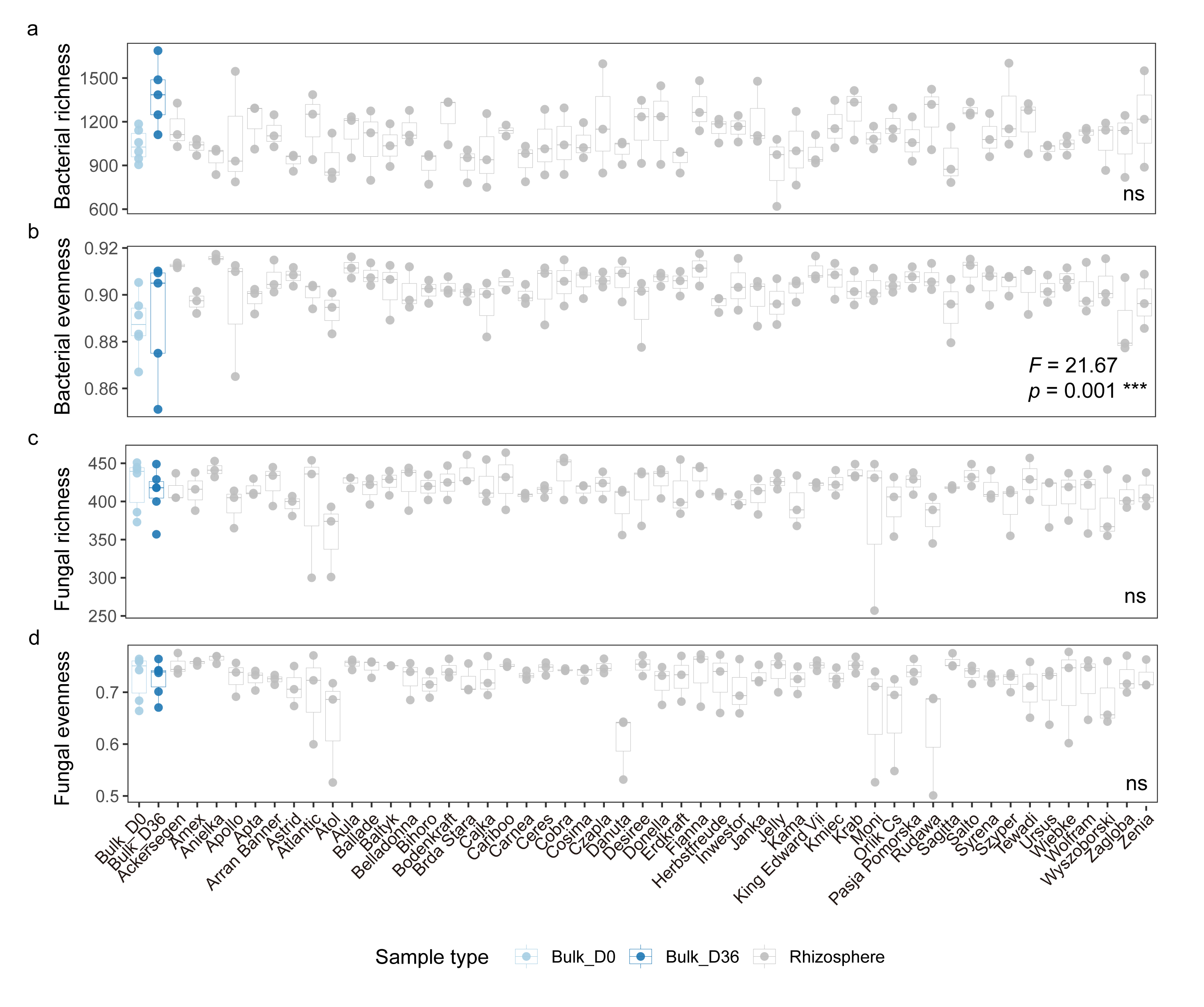
**Figure S5.** The alpha-diversity of the bacterial (a,b) and fungal (c,d) communities in bulk and rhizosphere soil, displayed by species richness and evenness. Different colours represent bulk samples from the beginning of the experiment (Bulk_D0), at harvest (Bulk_D36), and rhizosphere samples of 51 cultivars. The lower right corner of each plot displays one-way ANOVA results, where the F-value explains the variation among different soil compartments (bulk and rhizosphere), and the *p*-value indicates the statistical relationship. The symbol *** denotes statistically significant *p*-values (*p* = 0.001). The “ns” indicates no significant effect of soil compartments.

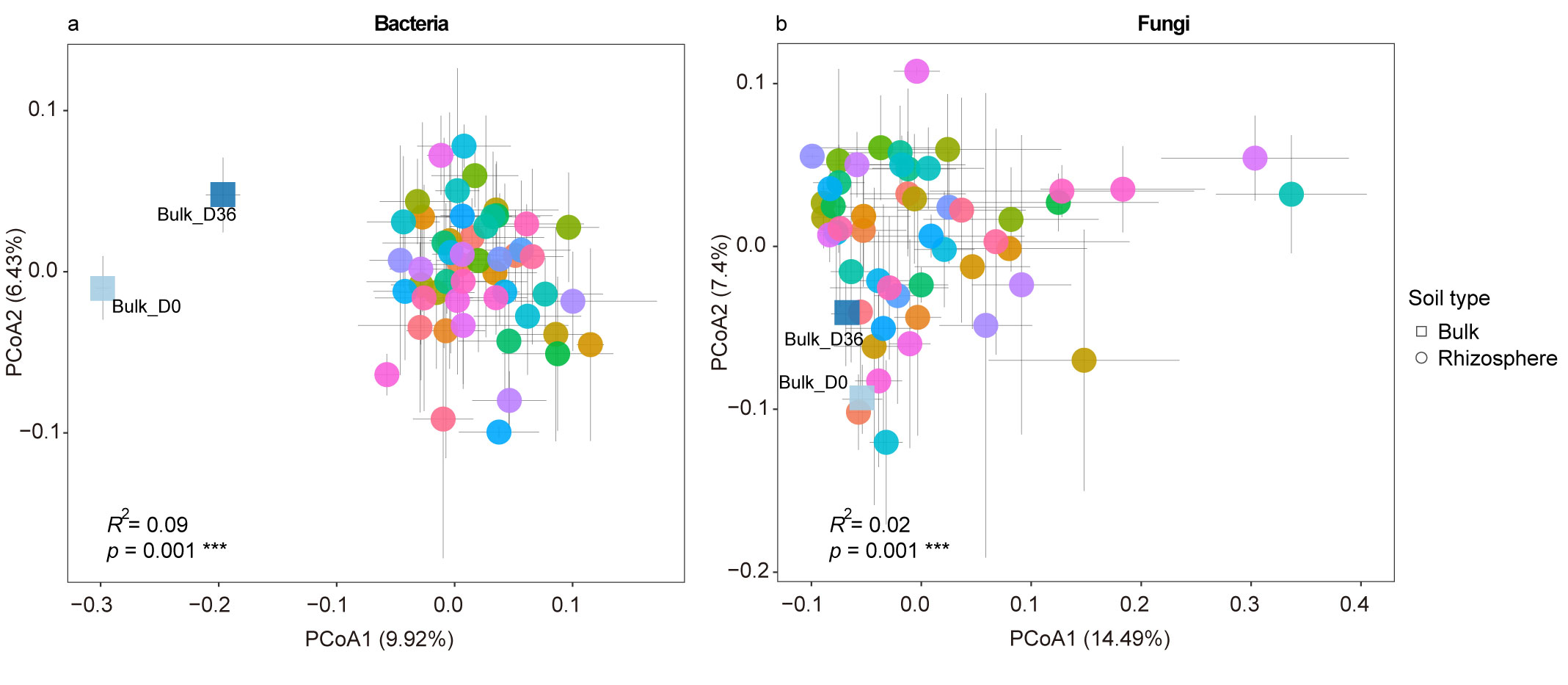
**Figure S6.** The microbial community composition in bulk and rhizosphere soil. Principal Coordinates Analysis (PCoA) based on Bray-Curtis dissimilarity was performed to visualise the community composition of bacteria (a) and fungi (b). Different shapes represent different soil compartments (bulk and rhizosphere). Different colours represent bulk samples from the beginning of the experiment (Bulk_D0), at harvest (Bulk_D36), and different potato cultivars, with the error bars for each cultivar displayed in grey. PERMANOVA results in the lower left corner of each panel elucidate the influence of soil compartments on community composition. *R*² quantifies the explained variation, and *p*-values are derived from 9999 permutations. The symbol *** denotes statistically significant *p*-values (*p* = 0.001).

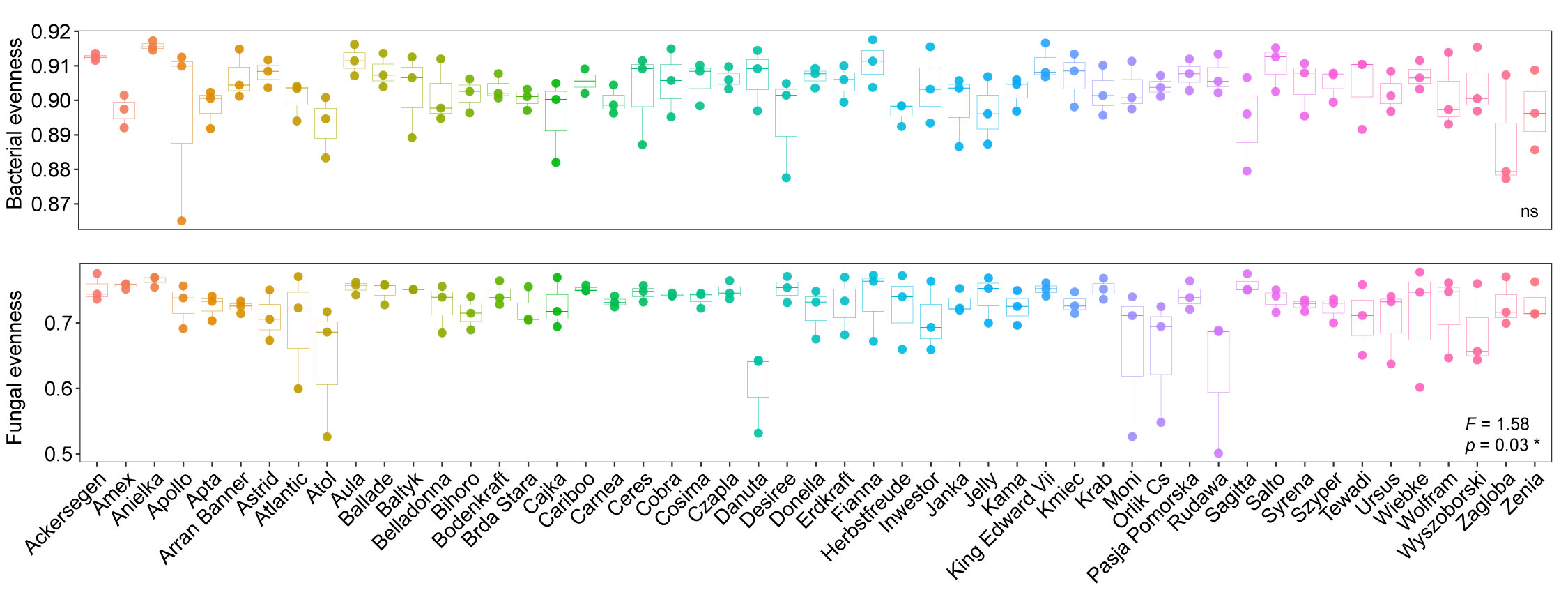
**Figure S7.** Rhizosphere microbial community evenness of 51 potato cultivars. The upper panel is bacterial evenness, and the lower panel is fungal evenness. Different colours indicate different cultivars. The lower right corner of each plot displays one-way ANOVA results, where the F-value explains the variation among different cultivars, and the *p*-value indicates the statistical relationship among cultivars. The symbol * denotes statistically significant p-values (*p* < 0.05). The “ns” indicates no significant effect. The microbial community species richness and Shannon diversities showed no significant difference among the 51 potato cultivars (data not shown).

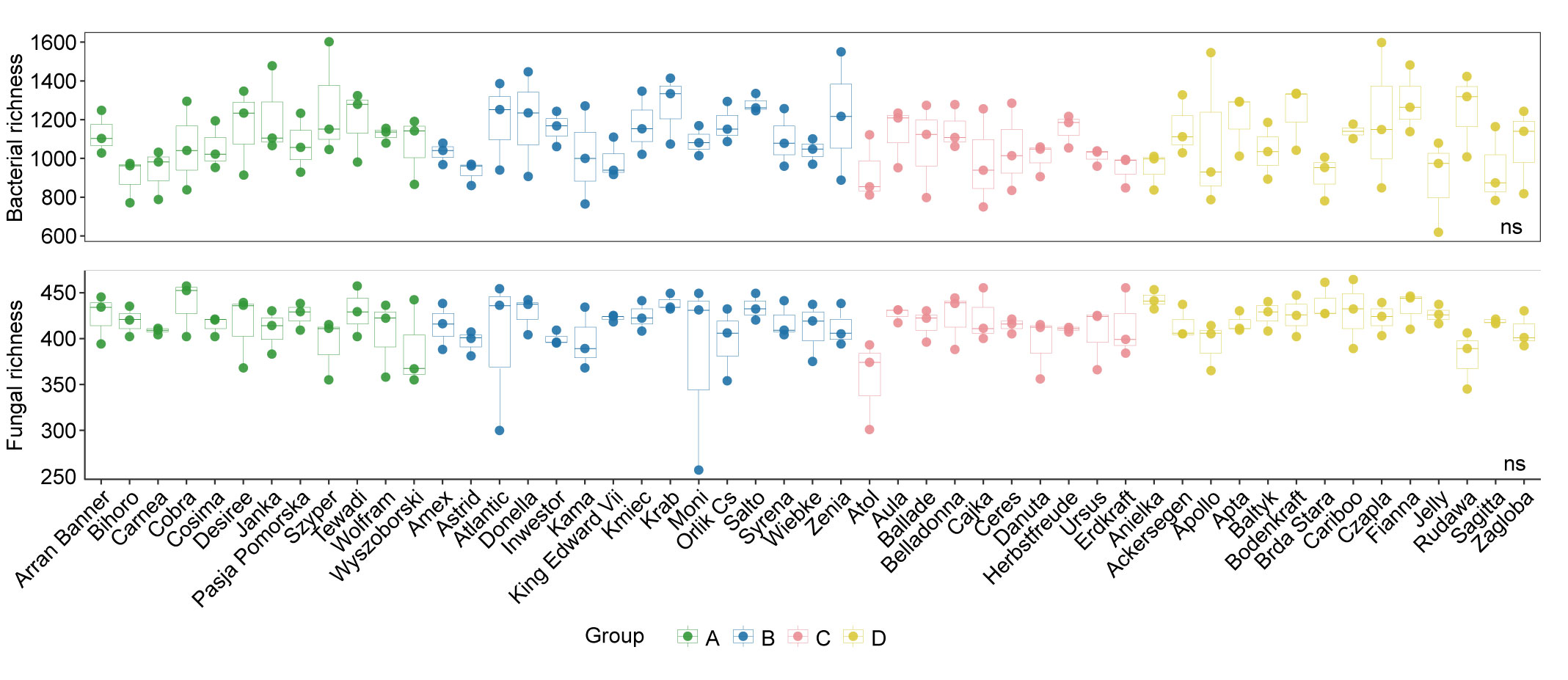
**Figure S8.** Rhizosphere microbial community species richness. The upper panel is bacterial richness, and the lower panel is fungal richness. Different colours indicate different cultivar functional groups. The “ns” indicates no significant influence of groups on microbial alpha diversity (one-way ANOVA). The microbial community evenness and Shannon diversities showed no significant difference among the functional groups (data not shown).

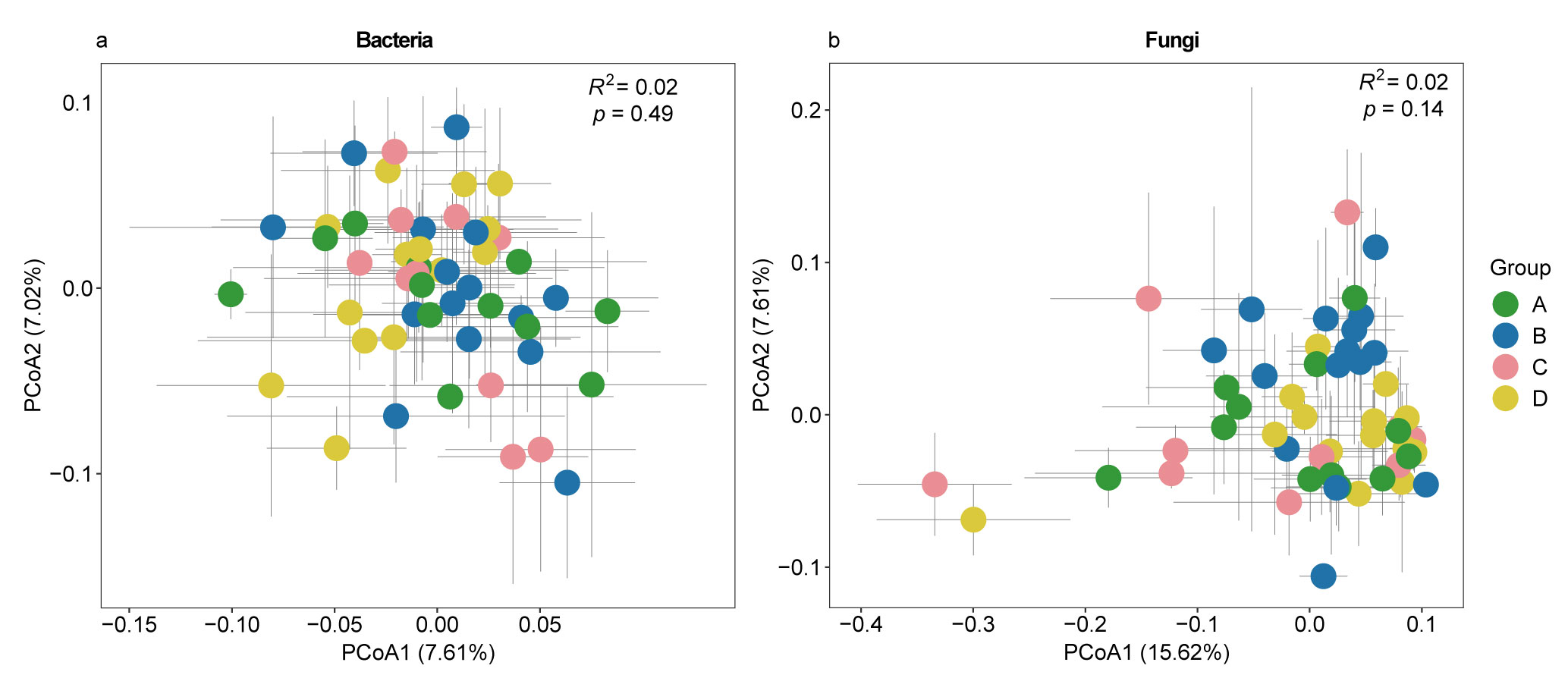
**Figure S9.** Rhizosphere microbial community composition. Principal Coordinates Analysis (PCoA) based on Bray-Curtis dissimilarity was performed to visualise the community dissimilarities of bacteria (a) and fungi (b). Distinct potato cultivar functional groups are represented by different colours, with the error bars for each cultivar displayed in grey. PERMANOVA results in the upper right corner of each panel elucidate the influence of groups on community composition. *R*² quantifies the explained variation, and *p*-values are derived from 9999 permutations.

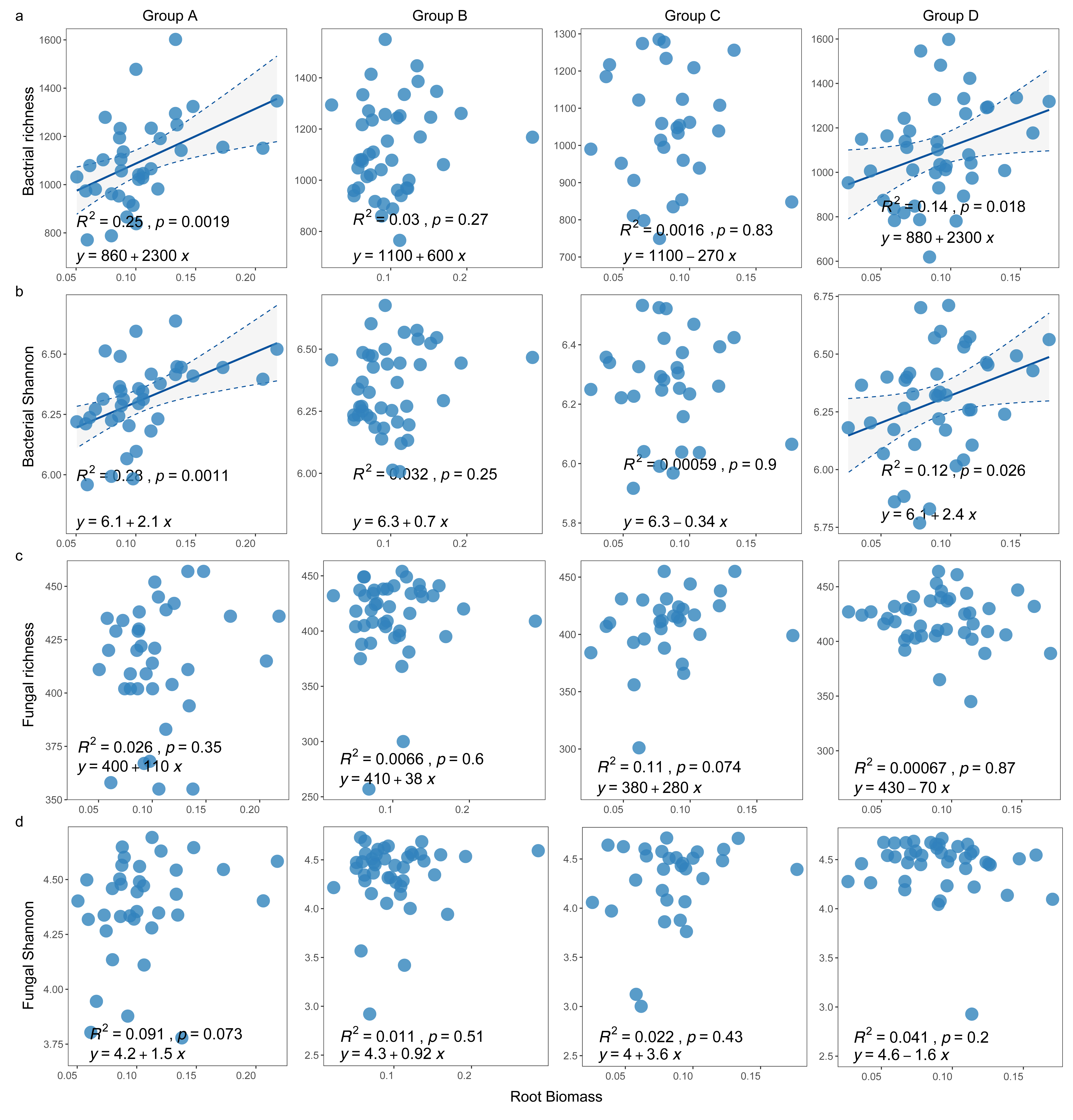
**Figure S10.** Correlation between plant root biomass and rhizosphere bacterial (a,b) and fungal (c,d)alpha diversities (species richness and Shannon diversity) across different functional groups. Displayed by functional groups A, B, C and D. The Spearman correlation assessed the relationship, with *R*^2^ indicating the strength of the correlation. Y is the regression equation, and a *p*-value < 0.05 represents a significant correlation between the variables.

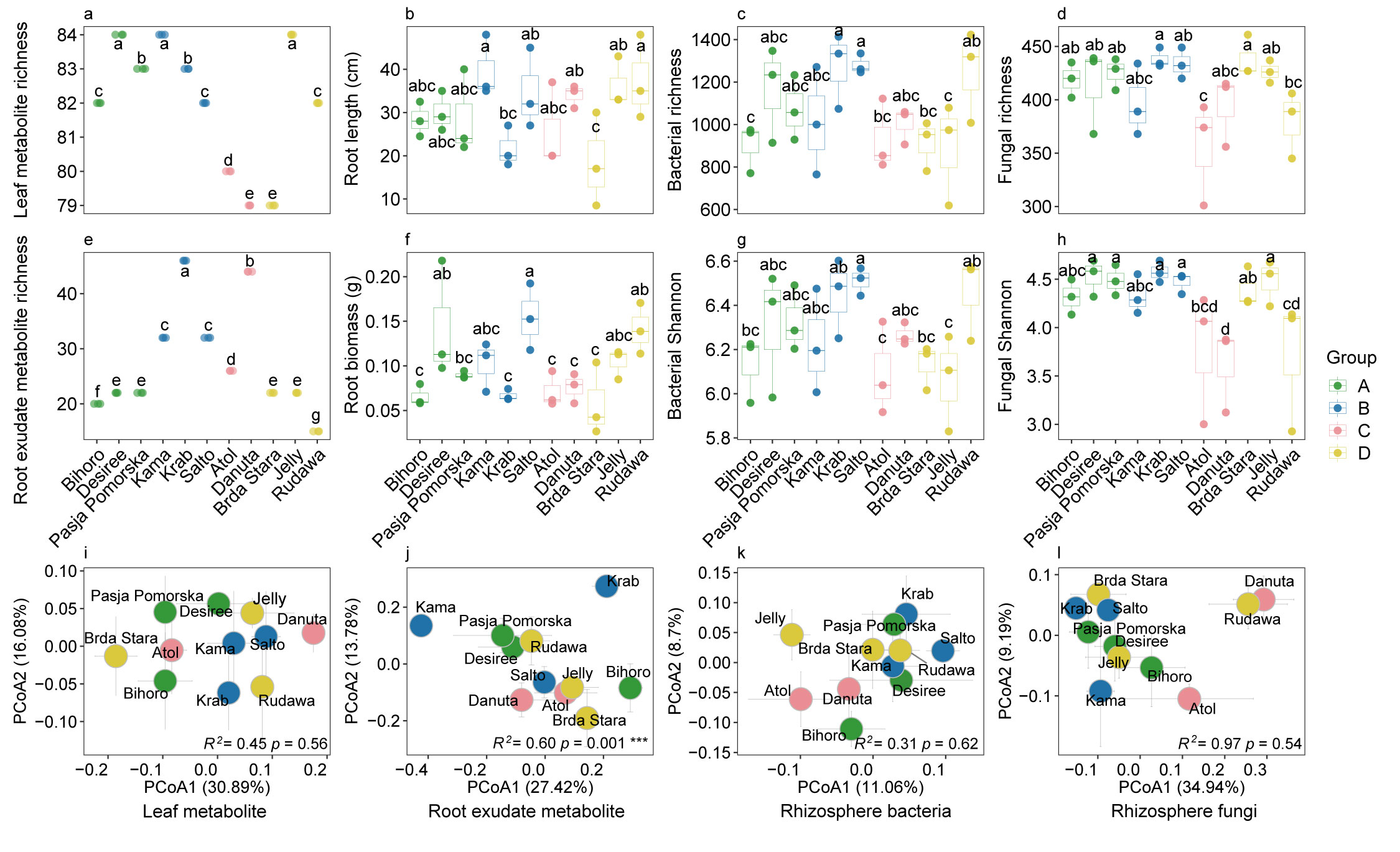
**Figure 11.** Characterisation of selected cultivars. Selected potato cultivars from different functional groups are represented by different colours, with the error bars for each cultivar displayed in grey. Letters in the upper two panels indicate significant differences across cultivars (Duncan post hoc test). In the lower right corner of the last panel plots, PERMANOVA results elucidate the influence of cultivars on community composition. *R*² quantifies the explained variation, and p-values are derived from 9999 permutations. A significance level is denoted as *** (*p* < 0.001).

**Table S1.** Background information on 148 selected cultivars for the *in* *vitro* experiment. "Y" indicates yes, "N" indicates no, and "NA" indicates no information available. Resistance is shown on a log10 scale ranging from 1 to 9, where 1 represents a lack of resistance, and 9 represents extreme resistance.

Table S1 can be found via the link: https://github.com/tianci-zhao/potatoMETAbiome-Greenhouse-Experiment.

**Table S2.** Plant growth data of 148 potato cultivars from *in vitro* experiment.

Table S2 can be found via the link: https://github.com/tianci-zhao/potatoMETAbiome-Greenhouse-Experiment.

**Table S3.** Soil physicochemical characteristics at the beginning and end of greenhouse experiment.

| Time point | Replicate | Organic Matter % | NO₃⁻ mg/kg | NH₄⁺ mg/kg | C:N | Moisture % | pH (H_2_O) | pH (KCl) |
| --- | --- | --- | --- | --- | --- | --- | --- | --- |
| beginning | 1 | 3.39 | 3.64 | 3.22 | 13.76 | 12.14 | 5.9 | 5.2 |
| beginning | 2 | 3.27 | 3.98 | 3.73 | 13.99 | 11.94 | 6.05 | 5.2 |
| beginning | 3 | 3.29 | 3.47 | 2.20 | 15.93 | 10.91 | 5.85 | 5.05 |
| beginning | 4 | 3.49 | 3.74 | 5.28 | 13.85 | 11.91 | 5.9 | 5.15 |
| beginning | 5 | 3.25 | 3.91 | 1.90 | 14.20 | 11.33 | 5.6 | 4.85 |
| beginning | 6 | 3.40 | 4.08 | 3.76 | 14.49 | 12.14 | 5.7 | 4.95 |
| end | 1 | 3.26 | 1.04 | 2.46 | 15.14 | 13.84 | 6.1 | 5.1 |
| end | 2 | 3.23 | 1.47 | 2.56 | 14.18 | 13.75 | 6 | 5.1 |
| end | 3 | 3.45 | 0.73 | 2.68 | 13.70 | 11.44 | 6.05 | 5.1 |
| end | 4 | 3.38 | 0.74 | 2.73 | 14.22 | 11.21 | 6.05 | 5.1 |
| end | 5 | 3.32 | 0.71 | 2.65 | 14.08 | 10.95 | 6.1 | 5.1 |
| end | 6 | 3.32 | 1.53 | 2.93 | 13.96 | 11.13 | 6.1 | 5.1 |

**Table S4.** Functional groups of potato cultivars categorised based on plant growth and metabolite profiles.

| Functional groups | Traits | Cultivars |
| --- | --- | --- |
| A | High leaf metabolite diversity and plant biomass | Arran Banner, Bihoro*, Carnea, Cobra, Cosima, Desiree*, Janka, Pasja Pomorska*, Szyper, Tewadi, Wolfram, Wyszoborski |
| B | High root exudates and plant length | Amex, Astrid, Atlantic, Donella, Inwestor, Kama*, King Edward VII, Kmiec, Krab*, Moni, Orlik Cs, Salto*, Syrena, Wiebke, Zenia |
| C | Complex leaf metabolite composition | Atol*, Aula, Ballade, Belladonna, Cajka, Ceres, Danuta*, Herbstfreude, Ursus, Erdkraft |
| D | Low root exudate metabolite diversity and different metabolite compositions | Ackersegen, Apollo, Apta, Anielka, Baltyk, Bodenkraft, Brda Stara*, Cariboo, Czapla, Fianna, Jelly*, Rudawa*, Sagitta, Zagloba |

*Indicates that these cultivars were suggested for future studies

**Table S5.** The one-way analysis of variance (ANOVA) shows the influence of cultivar on the plant metabolite alpha diversity. Significant *p*-values (*p* < 0.05) are shown in bold.

| Metabolite | Diversity | Df | F | *p*-value |
| --- | --- | --- | --- | --- |
| Leaf tissue | Richness | 50 | 1.54E+29 | **<0.001** |
|  | Evenness | 50 | 1.49 | **<0.05** |
|  | Shannon | 50 | 2.48 | **<0.001** |
| Root exudate | Richness | 50 | 22.98 | **<0.001** |
|  | Evenness | 50 | 1.9 | **<0.01** |
|  | Shannon | 50 | 6.27 | **<0.001** |

**Table S6.** Microbial interactive traits (MIT) levels of 51 potato cultivars.

| MIT levels | Cultivars |
| --- | --- |
| High | Ballade, Amex, Atlantic, Inwestor, Pasja Pomorska*, Tewadi, Rudawa*, Bodenkraft, Cajka, Desiree*, Janka, Kmiec, Czapla, Kama*, Fianna, Salto* |
| Middle | Danuta*, Wolfram, King Edward Vii, Krab*, Anielka, Apta, Zenia, Donella, Erdkraft, Carnea, Cobra, Syrena, Aula, Jelly*, Szyper, Arran Banner, Ursus |
| Low | Orlik Cs, Atol*, Moni, Bihoro*, Apollo, Wiebke, Ackersegen, Herbstfreude, Ceres, Brda Stara*, Cariboo, Cosima, Baltyk, Sagitta, Zagloba, Astrid, Wyszoborski |

*Indicates that these cultivars were suggested for future studies

**Data S1.** Dissolved organic carbon content of root exudates from in vitro experiment

Data S1 can be found via the link: https://github.com/tianci-zhao/potatoMETAbiome-Greenhouse-Experiment.

**Data S2.** Leaf tissue metabolites data in greenhouse experiment

Data S2 can be found via the link: https://github.com/tianci-zhao/potatoMETAbiome-Greenhouse-Experiment.

**Data S3.** Root exudate metabolites data *in vitro* experiment

Data S3 can be found via the link: https://github.com/tianci-zhao/potatoMETAbiome-Greenhouse-Experiment.

**Data S4.** Plant performance in greenhouse experiment

Data S4 can be found via the link: https://github.com/tianci-zhao/potatoMETAbiome-Greenhouse-Experiment.

**Data S5.** Rhizosphere microbial feature tables

Data S5 can be found via the link: https://github.com/tianci-zhao/potatoMETAbiome-Greenhouse-Experiment.

**Data S6.** MIT z scores data

Data S6 can be found via the link: https://github.com/tianci-zhao/potatoMETAbiome-Greenhouse-Experiment.
